# GDSL-domain containing proteins mediate suberin biosynthesis and degradation, enabling developmental plasticity of the endodermis during lateral root emergence

**DOI:** 10.1101/2020.06.25.171389

**Authors:** Robertas Ursache, Cristovao De Jesus Vieira-Teixeira, Valérie Dénervaud Tendon, Kay Gully, Damien De Bellis, Emanuel Schmid-Siegert, Tonni Grube Andersen, Vinay Shekhar, Sandra Calderon, Sylvain Pradervand, Christiane Nawrath, Niko Geldner, Joop E.M. Vermeer

## Abstract

Roots anchor plants and deliver water and nutrients from the soil. The root endodermis provides the crucial extracellular diffusion barrier by setting up a supracellular network of lignified cell walls, called Casparian strips, supported by a subsequent formation of suberin lamellae. Whereas lignification is thought to be irreversible, formation of suberin lamellae was demonstrated to be dynamic, facilitating adaptation to different soil conditions. Plants shape their root system through the regulated formation of lateral roots emerging from within the endodermis, requiring local breaking and re-sealing of the endodermal diffusion barriers. Here, we show that differentiated endodermal cells have a distinct auxin-mediated transcriptional response that regulates cell wall remodelling. Based on this data set we identify a set of GDSL-lipases that are essential for suberin formation. Moreover, we find that another set of GDSL-lipases mediates suberin degradation, which enables the developmental plasticity of the endodermis required for normal lateral root emergence.

## INTRODUCTION

Plants require a dynamic root system for optimal anchorage and to forage the soil environment for water and nutrients. Therefore, both functions need to be fine-tuned for optimal performance under challenging and fluctuating growth conditions. The root system architecture is modulated by regulated branching through the formation of lateral roots. In most angiosperms, including Arabidopsis thaliana (Arabidopsis), these organs initiate in the pericycle from de novo-formed meristems, often adjacent to the xylem, called xylem-pole pericycle cells (XPP). Auxin is required both for the initiation as well as for the development of lateral roots (Banda et al., 2019; Stoeckle et al., 2018). In Arabidopsis, lateral roots need to traverse three overlying cell layers in order to emerge; the endodermis, cortex and epidermis. Since plant cells are interconnected through their cell walls and under considerable turgor pressure, the newly formed organ must overcome the mechanical constraints posed by these cell layers. Especially the direct neighbor of the XPP, the endodermis, plays an essential role during lateral root formation, as it actively accommodates the expansion growth of the XPP through remodeling of cell shape and volume. It was shown that these responses were regulated via auxin-mediated signaling, and endodermal expression of a dominant repressor of auxin signaling, short hypocotyl 2-2 (shy2-2), completely blocked lateral root formation. Thus, after auxin-mediated priming, lateral root founder cells swell and this expansion growth needs to be accommodated via SHY2-mediated endodermal auxin signaling (Vermeer et al., 2014). The observation that lateral root primordia grow through the overlying endodermal cells without compromising their viability, is in agreement that this process is associated with extensive local cell wall remodeling (Kumpf et al., 2013; Swarup et al., 2008). In addition, it was reported that the Casparian Strip (CS), a lignified primary cell wall modification acting as a diffusion barrier, appears to get locally degraded in order to allow the growth of the lateral root through this cell layer (Li et al., 2017). It was also shown that suberin, a secondary cell wall modification, is deposited in the cell walls of endodermal cells in contact with the lateral root primordium during emergence. However, endodermal cells often are already suberized when lateral root formation occurs and it is still unknown how suberin is first degraded and later re-synthesized. The growth of a lateral root through the endodermis needs to be tightly-controlled, in order to minimize both the leakage of nutrients from the stele into the rhizosphere and the entry of soil-borne pathogens to enter the stele. Therefore, a dynamic de-suberization and re-suberization is bound to play an important role in this process. It has been demonstrated that ABA- and ethylene signaling pathways are crucial for nutrient-induced plasticity of endodermal suberization during plant development (Barberon et al., 2016). However, although several studies have revealed roles of suberin deposition in adaptation to the soil environment and organ formation, we still lack understanding of the basic molecular machineries regulating suberin biosynthesis and deposition during root development (Andersen et al., 2020; Barberon et al., 2016; Li et al., 2017; Yadav et al., 2014). This is especially relevant for understanding the localized, dynamic suberin deposition found in roots. In this study we used the *CASP1pro::shy2-2* line as a tool to identify cell wall remodeling factors during this process. The *CASP1pro::shy2-2* mutant is specifically blocked in endodermal auxin-responses, which strongly impairs lateral root formation. In order to obtain a specific, endodermal auxin-response profile, we used a combination of *CASP1pro::shy2-2* and s*olitary root 1* (*slr-1*), two mutants that repress auxin signaling in different spatial domains (Fukaki et al., 2002; Swarup et al., 2008; Vermeer et al., 2014). In the obtained data set, we identify a distinct set of 10 GDSL-motif containing enzymes that are differentially expressed between *slr-1* and *CASP1pro::shy2-2* after auxin treatment. All of these ten GDSL-motif containing enzymes were either expressed in the endodermis, repressed or induced in this cell layer during auxin treatment or lateral root formation. We show that five of the GDSL-motif containing enzymes that are repressed by auxin are required for suberin biosynthesis, whereas another five auxin-induced GDSL-motif containing enzymes are required for suberin degradation in the endodermis during lateral root formation. Quintuple mutants of the suberin biosynthesis GDSL-motif containing enzymes were overly-sensitive to mild salt stress, which resulted in reduced fresh weight and a strong reduction in emerged lateral roots. Single knock-out mutants of several members of the auxin-induced group of suberin degrading enzymes resulted in a delayed lateral root emergence. Our work reveals for the first time essential enzymatic components that regulate suberin polymerization and degradation, strongly impacting our understanding of in vivo suberin formation, as well as its striking developmental plasticity.

## RESULTS

### SHY2-mediated transcriptional responses in differentiated endodermal cells

We have previously shown that during lateral root formation in Arabidopsis, overlying endodermal cells undergo drastic changes in cell volume and that the CS, a lignified primary cell wall modification, is locally modified to facilitate lateral root emergence (Vermeer et al., 2014). Furthermore, we showed that *SHORT HYPOCOTYL 2* (*SHY2/IAA3*)-mediated auxin signaling drives these responses. SHY2 represses its own transcription in a typical, auxin-induced negative feedback loop and is thus a great, early transcriptional auxin-response marker (Tian et al., 2002; Vermeer et al., 2014). However, we do not know which SHY2 targets (direct or indirect) are required for the complex accommodating responses in the endodermis. Therefore, we set out to obtain a SHY2-mediated transcriptional response profile in the endodermis. Generating such a data set comes with particular challenges. First, most endodermal cells at the moment of lateral root emergence are lignified and suberized, making it impossible to employ protoplast isolation used for single cell or cell-type specific sequencing. Secondly, only endodermal cells overlying an auxin-emitting lateral root primordium from stage I and onwards will be stimulated in a SHY2-dependent fashion (Vermeer et al., 2014). Thus, only a subset of endodermal cells would display the transcriptome profile we are interested in. We therefore first thought to compare wild-type and *CASP1pro::shy2-2* seedlings treated with auxin to obtain a differential, endodermis-specific set of auxin responsive genes. In wild-type, *SHY2pro::NLS-3xmVENUS* fluorescence peaks in the endodermis at ∼16 hours after NAA treatment (Figure 1A and D). However, since the *CASP1pro::shy2-2* mutant impairs the auxin-mediated induction of lateral root formation that is strongly induced in wild-type (Vermeer et al., 2014), a direct comparison of the auxin-induced transcriptomes of *CASP1pro::shy2-2* roots to wild-type roots would result in a strong bias for pericycle and cell cycle-related genes and not necessary genes involved in endodermal accommodating responses. Therefore, we designed a genetic trick to enrich for auxin induced transcriptional changes in the endodermis.

**Figure 1.**
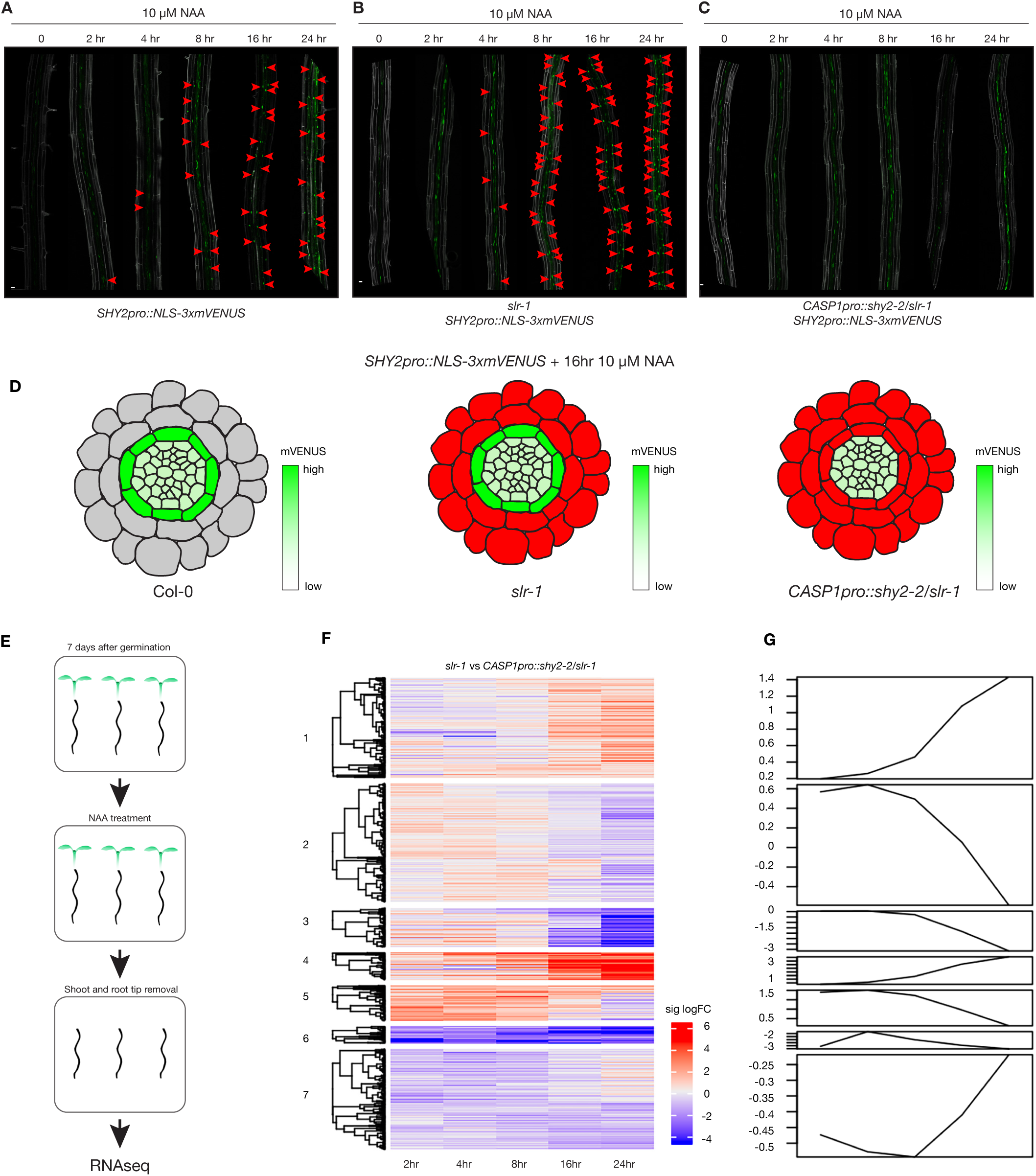
A genetic trick for mapping auxin responses in the differentiated endodermis. **A-C**. Maximum images projections of roots expressing *SHY2pro::NLS-3xmVENUS* treated with 10 µM NAA for 0, 2, 4, 8, 12 and 24hrs. **A**. Col-0. **B**. *slr-1*. **C**. *CASP1pro::shy2-2/slr-1*. Red arrowheads indicate *SHY2pro::NLS-3xmVENUS* signal in the endodermis. **D**. Schematic representation of *SHY2pro::NLS-3xmVENUS* responses in the different genetic backgrounds after 10 µM NAA for 16hrs. Red indicates block of auxin signalling, green indicates induction of *SHY2* **E**. Experimental setup of the RNAseq experiment. **F**. Heatmap showing the seven clusters containing the significant differentially expressed genes between *slr-1* and *CASP1pro::shy2-2/ slr-1* during the NAA time course. **G**. Graphical presentation of the behavior of the seven cluster during the NAA time course depicted in (**E**). Scale bar in (**A**) = 50 µm.

To this end, we combined the dominant, *solitary root 1* (*slr-1*)/*IAA14* mutant with *CASP1pro::shy2-2*. Lateral root formation is impaired in the Arabidopsis *slr-1* mutant (Fukaki et al., 2002). Importantly, *SLR* is expressed in the pericycle, cortex and epidermis, but not in the endodermis and the slr-1 mutant should thus specifically block auxin response in the surrounding cell layers (Fukaki et al., 2002; Swarup et al., 2008)(Figure S1). Indeed, we found that auxin-mediated induction of *SHY2* in the endodermis was still occurring in *slr-1* roots (Figure 1B and D). We predicted that, in the combined *CASP1pro::shy2-2* /*slr-1* background auxin signaling is basically blocked in all differentiated root cell layers. As mentioned above, SHY2 represses its own expression (Tian et al., 2002), and indeed, as shown in Figure 1C, we could not detect induction of *SHY2pro::NLS3xmVENUS* in the endodermis of the *CASP1pro::shy2-2*/*slr-1* background. Based on these results, we predicted that a comparison (subtraction) of the NAA-induced transcriptomes of roots from the *slr-1* single mutant with the *CASP1pro::shy2-2*/*slr-1* double mutant would allow us to extract a specific endodermal auxin signaling transcriptomic profile, otherwise obscured by the strong, proliferation-inducing auxin responses of the xylem-pole pericycle cells (Figure 1A-E).

### Differentiated endodermal cells have a distinct auxin-mediated transcriptional response

We interrogated the genome-wide transcriptional responses in *slr-1* and *CASP1pro::shy2-2*/*slr-1* after NAA treatment (10µM) at multiple time points. After statistical analysis (fold change > 2, false discovery rate < 0.05), we established that ∼800-900 genes are differentially expressed at 2, 4, 8, 16hr after treatment and ∼1000 are significantly changed after 24hr treatment compared to the zero timepoint (Table S1). Using non-supervised methods and manual tests we settled on 7 clusters to describe the data (Figure 1F and G). As expected, the data set contained a large number of cell wall-related genes and hardly any cell cycle-related genes. When looking at the gene ontology (GO) annotations, we observed terms linked to auxin signaling and lateral root development (cluster 2 and 5), whereas terms related to lipid transport and fatty acid metabolism were enriched in clusters 3 and 5. The fact that we observed in general little GO terms related solely to auxin signaling and lateral root development is most likely due to the unique experimental design, providing a previously undescribed auxin-response profile focused on a specific, differentiated cell type. To substantiate this, we compared the *slr-1* versus *CASP1pro::shy2-2*/*slr-1* data with the two other published data sets from transcriptome analysis dealing either with roots treated with auxin or microdissection of root sections after gravistimulation mediated of lateral root induction (Lewis et al., 2013; Voß et al., 2015). After re-analyzing the differential gene-expression analysis for both data sets, we converted the logFC values of each gene to z-scores in order to facilitate the comparison of the different data sets. Interestingly, there appeared to be little correlation between the differentially expressed genes in our data set and those in the data sets of Lewis et al., (2013) or Voß et al., (2015) (Figure S2). We took this as a confirmation of the unique and specific nature of our transcriptional profile. In order to try and identify novel genes involved in cell wall modification, as well as to confirm the validity of our transcriptional profile, we selected a wide range of genes possibly related to the observed endodermal responses, including genes linked to lignification, lipid transport and as well as several unknown genes showing particularly strong and high-confidence differential responses. We generated promoter-reporter lines to characterize their expression pattern during root development and lateral root formation. In a strong validation of our approach, most of the selected genes were found to display auxin-regulated expression in the endodermis (Figure S3 and Table S2). The transcriptional reporter for one of such candidate, the GDSL-type esterase/lipase 12 (GELP12), was expressed in the cortex of wild-type and *CASP1pro::shy2-2* roots, but was not detected in the cortex of *slr-1* roots (Figure S3A). Treatment of *slr-1*/*GELP12pro::NLS-3xmVENUS* roots with auxin resulted in induction of *GELP12* specifically in the endodermis, but did not restore cortex expression, as auxin-mediated gene expression is blocked by the presence of dominant *slr-1* allele. Supplemental Figure S3C shows a selection of the candidates that all display expression in the endodermis (either constitutive or induced during lateral root formation). In contrast, the expression domain of *LACCASE 2* (*LAC2*), although showing significant differential expression between *slr-1* and *CASP1pro::shy2-2/ slr-1*, was not expressed in the endodermis, but rather in protoxylem cells (Figure S3D and Table S2). Nevertheless, out of the 27 candidates for which we generated transcriptional reporters, only 3 turned out not to be expressed or induced in the endodermis (Table S2). Since we were interested in possible cell wall modifying enzymes, we searched the list of differential expressed genes for cell wall-associated functions. This lead to the observation that many genes with functions attributed to cutin/suberin homeostasis such as *3-KETOACYL-COA SYNTHASE* (*KCS*), *FATTY ACID DESATURASE* (*FAD*), *FATTY ACID REDUCTASE* (*FAR*), *LIPID TRANSFER PROTEIN* (*LTP*), *GDSL-TYPE ESTERASE/LIPASE* (*GELP*) and *ATP-BINDING CASSETTE G* (*ABCG*) transporters (Edqvist et al., 2018; Fich et al., 2016; Lai et al., 2017; Salminen et al., 2018; Vishwanath et al., 2015) showed highly dynamic, differential expression in our dataset (Figure S3E). Suberin deposition has been shown to be highly plastic, might be continuously turned over and is also implicated in adaptation to the soil environment and during lateral root development (Andersen et al., 2020; Barberon et al., 2016; Li et al., 2017). However, fundamental aspects of suberin formation and deposition are still not understood, especially its synthesis, deposition and turnover in the apoplast. Therefore, we decided to investigate whether some of the cell wall-related differentially expressed genes in our endodermis-focused auxin response data set could be involved in suberization and reveal new insights into this process. In particular, we were intrigued by the high number of differentially expressed GELPs in our dataset (Figure S3E), since members of this large family have been shown to be involved in cutin polymerization, but also to be able to degrade both cutin and suberin (Bakan and Marion, 2017; Girard et al., 2012; Naseer et al., 2012; Philippe et al., 2016; Yeats et al., 2012). Thus, we decided to focus on the differentially regulated set of GELPs in our data set.

### Suberin is degraded while the lateral root cap cuticle is established during lateral root formation

Fluorol Yellow (FY) staining of suberin and cutin is dynamic during lateral root formation. However, FY stains both suberin and cutin (Berhin et al., 2019) (Figure S7A), so it is not yet clear if, how and at which stage suberin might be degraded in the endodermis and replaced by a cutin-like structure in the emerging primordium. Therefore, to get a deeper insight into this process, we analyzed the dynamics of suberin and cutin during lateral root formation using transmission electron microscopy (TEM) (Figure 2). Analyzing stage II lateral root primordia, which usually form in the unsuberized zone, we could detect suberin lamellae only in the endodermal cell walls facing the lateral root primordia, but not in the cell wall of endodermal cells on the opposite side of the root (Figure 2A). Stage III primordia are found in the patchy suberized zone of the root and here we expectedly detected both suberized and non-suberized endodermal cells. At this stage we started to distinguish the onset of the lateral root cap cuticle formation, accompanied by the disappearance of the suberin in the cell walls of the endodermal cells overlying the primordium (Figure 2B). In stage IV primordia, which are usually found in the fully suberized zone, we could detect suberin deposition in all endodermal cells in the zone of no developing lateral roots. At this point it was difficult to observe any suberin in cell walls of endodermal cells facing the primordia. Indeed, it appeared that the suberin was degraded in coordination with the formation of the root cap cuticle (Figure 2C). In fully emerged lateral roots, we could only detect the lateral root cap cuticle, whereas endodermal cells not in contact with the primordium still maintained their suberin lamellae (Figure 2D). Thus, it is clear that suberin is gradually degraded in cell walls of endodermal cells surrounding the lateral root primordium while the lateral root primordium synthesizes its lateral root cap cuticle as a protective coating during emergence (Berhin et al., 2019).

**Figure 2.**
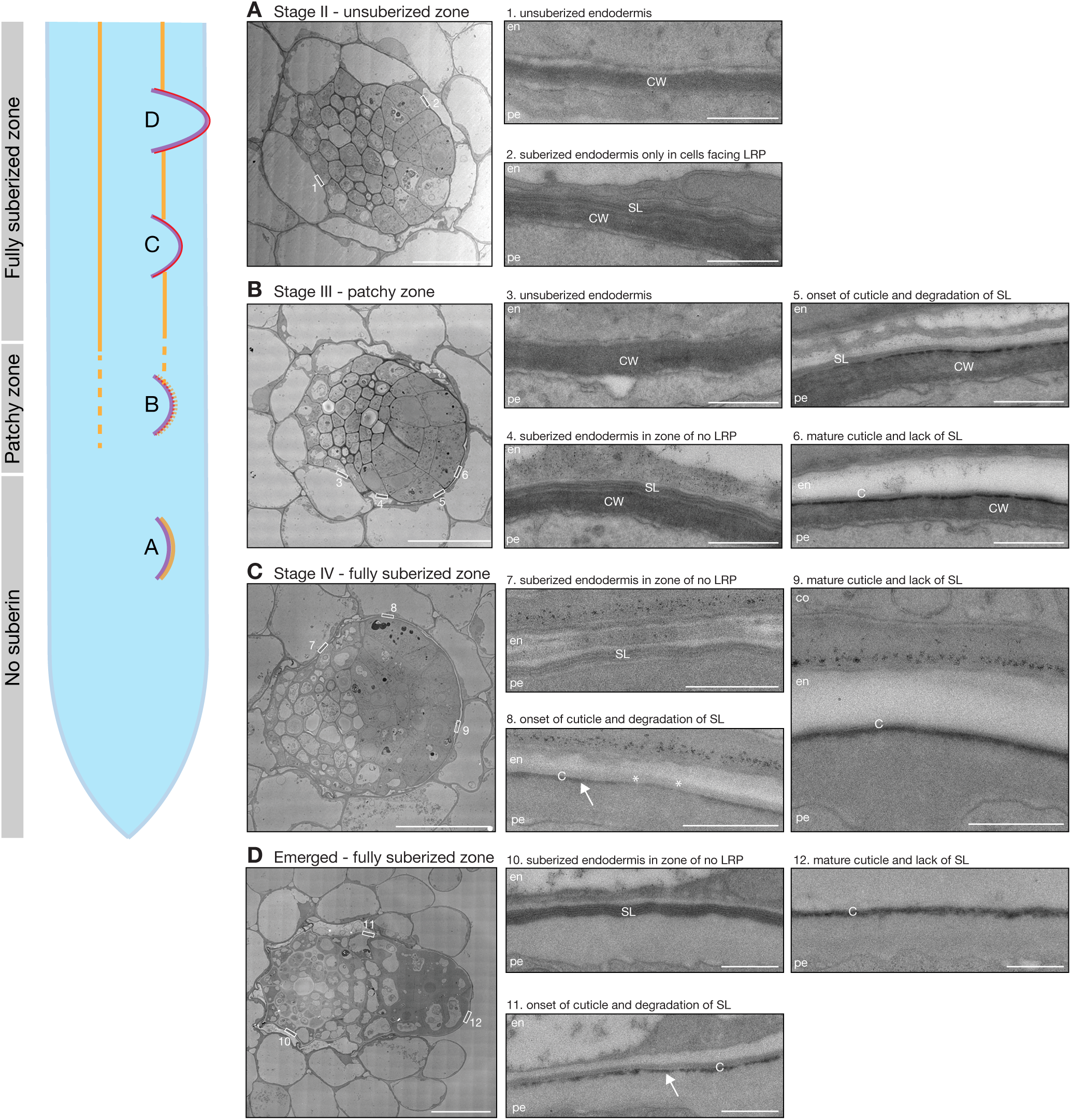
Suberin is degraded while the lateral root cap cuticle is established during lateral root formation. Schematic drawing of a root highlighting the different stages of suberin formation in the endodermis. Letters refer to the TEM images with the corresponding letter. Cell wall is shown in purple, suberin lamellae in yellow and the lateral root cap cuticle in red. **A**. TEM micrograph of a root section containing a stage II lateral root primordium. The numbered boxed regions are shown as magnifications marked by the same number. **B**. TEM micrograph of a root section containing a stage III lateral root primordium. The numbered boxed regions are shown as magnifications marked by the same number. **C**. TEM micrograph of a root section containing a stage IV lateral root primordium. The numbered boxed regions are shown as magnifications marked by the same number. **D**. TEM micrograph of a root section containing an emerged lateral root. The numbered boxed regions are shown as magnifications marked by the same number. En = endodermis, pe = pericycle, co = cortex, SL = suberin lamellae, C = lateral root cap cuticle, CW = cell wall. Scale bars in (**A**-**D**) = 10 µm and scale bars in (1-12) panels = 1 µm.

### Suberin deposition strongly requires a cluster of auxin-repressed GELPs

We have demonstrated that suberin deposition is regulated during lateral root formation and that it gradually is replaced by the lateral root cap cuticle (Figure 2). Whereas we have some insights regarding cutin polymerization within the apoplast, there are no factors known to mediate suberin polymerization (Philippe et al., 2020). Generally, very few strong mutants for suberin biosynthesis have been identified. *gpat5* mutants displayed reduced suberin levels in the main root and in the seed coat and were more susceptible to elevated salt concentrations. However, suberin deposition was only partially (∼30%) affected in *gpat5-1* mutants (Beisson et al., 2007). Currently, the strongest available interference with suberin in roots relies on either endodermis-specific interference with ABA or cytokinin signaling, or on artificial overexpression of a cutin-degrading enzyme (Andersen et al., 2018; Barberon et al., 2016; Naseer et al., 2012).

It was previously demonstrated that a member of the large family of GELP proteins (Yeats et al., 2012), *CUTIN DEFICIENT 1* (*CD1*), has cutin in vitro synthase activity and *CD1* loss-of-function mutants in tomato show partial defects in cuticle formation, but no equivalent evidence exists for suberin synthases. Interestingly, we observed a group of five GELPs (*GELP22, 38, 49, 51* and *96*) to be differentially downregulated after prolonged auxin treatment (Figure 3A). Since auxin treatment results in a massive induction of lateral root formation, accompanied by degradation of suberin in overlying endodermal cells (Li et al., 2017), we hypothesized that suberin biosynthetic enzymes would be inhibited during formation of lateral root primordia. Therefore, we speculated that the five GELPs repressed by auxin treatment might have a role in suberin biosynthesis. This idea was corroborated by their expression pattern in the root obtained with transcriptional reporters for *GELP22, 38, 49, 51* and *96*. Clearly, *GELPXpro::NLS-3xmVENUS* reporter lines revealed endodermis-specific expression for *GELP38, GELP51* and *GELP96*, and expression in endodermis and epidermis for *GELP22* and *GELP49* (Figure 3C). Since treatment of Arabidopsis seedlings with ABA results in a significant increase in suberin deposition and *GPAT5* marker expression (Barberon et al., 2016), we further checked whether *GELP22, 38, 49, 51* and *96* would be induced by ABA treatment. All *GELP* reporter lines were induced in response to ABA treatment and also expanded their expression domain into the cortex, very similar to what has been reported for *GPAT5* (Barberon et al., 2016) and (Figure S4A and B). Together, our data establishes a strong correlation between suberin biosynthesis and the expression pattern of these five GELPs. In order to establish a function for these GELPs in suberin biosynthesis, we collected available T-DNA insertion mutants and characterized these for differences in suberin deposition using FY or Nile Red staining, two fluorescent dyes that both stain suberin (Figure 3B). In the absence of T-DNA insertion lines for *GELP38*, we generated two loss-of-function mutants using CRISPR/Cas9. None of the single knock-out or knock-down mutants showed any significant difference in suberin occupancy in Arabidopsis roots compared to the wild-type control (Figure S4C-E). Because of their likely functional redundancy, we generated two different allelic combinations of the five putative suberin biosynthesis-related GELPs: *gelp22-c1*/*gelp38-c3*/*gelp49-c1*/*gelp51-c1*/*gelp96-c1* and *gelp22-c2*/*gelp38-c4*/*gelp49-c2*/*gelp51-c2*/*gelp96-c2* (hereafter called *gelp*^*quint-1*^ and *gelp*^*quint-2*^) (Figure S5). To test whether suberin levels in roots of the *gelp*^*quint-1*^ and *gelp*^*quint-2*^ mutants were affected compared to wild-type, we stained roots of 5-day old plants with FY and Nile Red. Whereas suberin staining in wild-type resulted in the same pattern as described before (Naseer et al., 2012), both quintuple mutants showed a complete absence of suberin staining (Figure 4A-D and Figure S6A). ABA treatment strongly enhances suberin deposition in both the endodermis and cortex (Barberon et al., 2016). Yet, despite this ABA-induced boost in suberin production, we again could not detect any suberin deposition using FY staining in the roots of *gelp*^*quint-1*^ and *gelp*^*quint-2*^ (Figure 4E and F). Since cutin and suberin are similar in structure and both can be stained by FY, we then investigated whether the lateral root cap cuticle was also affected in roots of the *gelp*^*quint-1*^ mutant, using FY staining of emerging lateral roots. FY still stained the dome of the emerging lateral root, suggesting that the gelpquint mutants are specifically affected in suberization and that the lateral root cap cuticle is still intact (Figure 4G). The TEM analysis confirmed that the cuticle is formed normally in the *gelp*^*quint*^ mutants (Figure S6C). Since endodermal suberin was shown to interfere with uptake from the apoplast, we used the fluorescein diacetate (FDA) penetration assay (Barberon et al., 2016) to test for the presence of functional suberin lamellae in the endodermis of *gelp*^*quint-1*^ and *gelp*^*quint-2*^ roots. Whereas FDA could only enter the epidermis and cortex in the suberized zone of wild-type roots, it was entering the endodermis of *gelp*^*quint-1*^ and *gelp*^*quint-2*^ roots, further supporting the absence or strong deficiency of a suberin barrier in the endodermis (Figure S6H-J). Both, the absence of FY staining in the *gelp*^*quint-1*^ and *gelp*^*quint-2*^ and the FDA uptake assay suggest that the *gelp*^*quint-1*^ and *gelp*^*quint-2*^ mutants have strongly reduced suberin deposition. To verify this, we performed a chemical analysis of the suberin content in roots of wild-type, *gelp*^*quint-1*^ and *gelp*^*quint-2*^ mutants. This revealed strong reductions in the amount of aliphatic and aromatic suberin monomers in *gelp*^*quint-1*^ and *gelp*^*quint-2*^ mutants that were nearly identical in both allelic combinations. Dicarboxylic acids were very nearly absent (98% reduction), while ω-hydroxy acids and fatty alcohols were reduced by ∼ 90% and 50%, respectively (Figure 4H). Ferulate was reduced by 70%, while coumarate showed only minor reductions. This resulted overall in a ∼85% reduction in the amount of suberin monomers compared to wild-type roots (Figure 4H), which correlates well with their FY phenotypes. Next, we complemented the suberin phenotype of the *gelp*^*quint-1*^ mutant by introducing a fluorescent protein-labelled biosynthetic GELP. Expression of an inducible *GELP38proXVE*>>*GELP38-mCitrine* fusion in the gelpquint-1 mutant restored the stereotypical FY staining in roots, indicating that the GELP38-mCitrine is a functional fusion protein. Moreover, we could show that GELP38-mCitrine is localized in the apoplast of the endodermis where suberin polymerization takes place (Figure 4I-K). We also tested whether the Casparian strip was unaffected in the *gelp*^*quint-1*^ and *gelp*^*quint-2*^ mutants. Both the lignin staining with Basic Fuchsin and uptake of Propidium Iodide (PI) were not significantly affected compared to wild-type (Figure S6D-G). Thus, a cluster of five auxin-repressed GELPs is essential for normal suberin deposition in the endodermis, but not for the formation of the lateral root cap cuticle.

**Figure 3.**
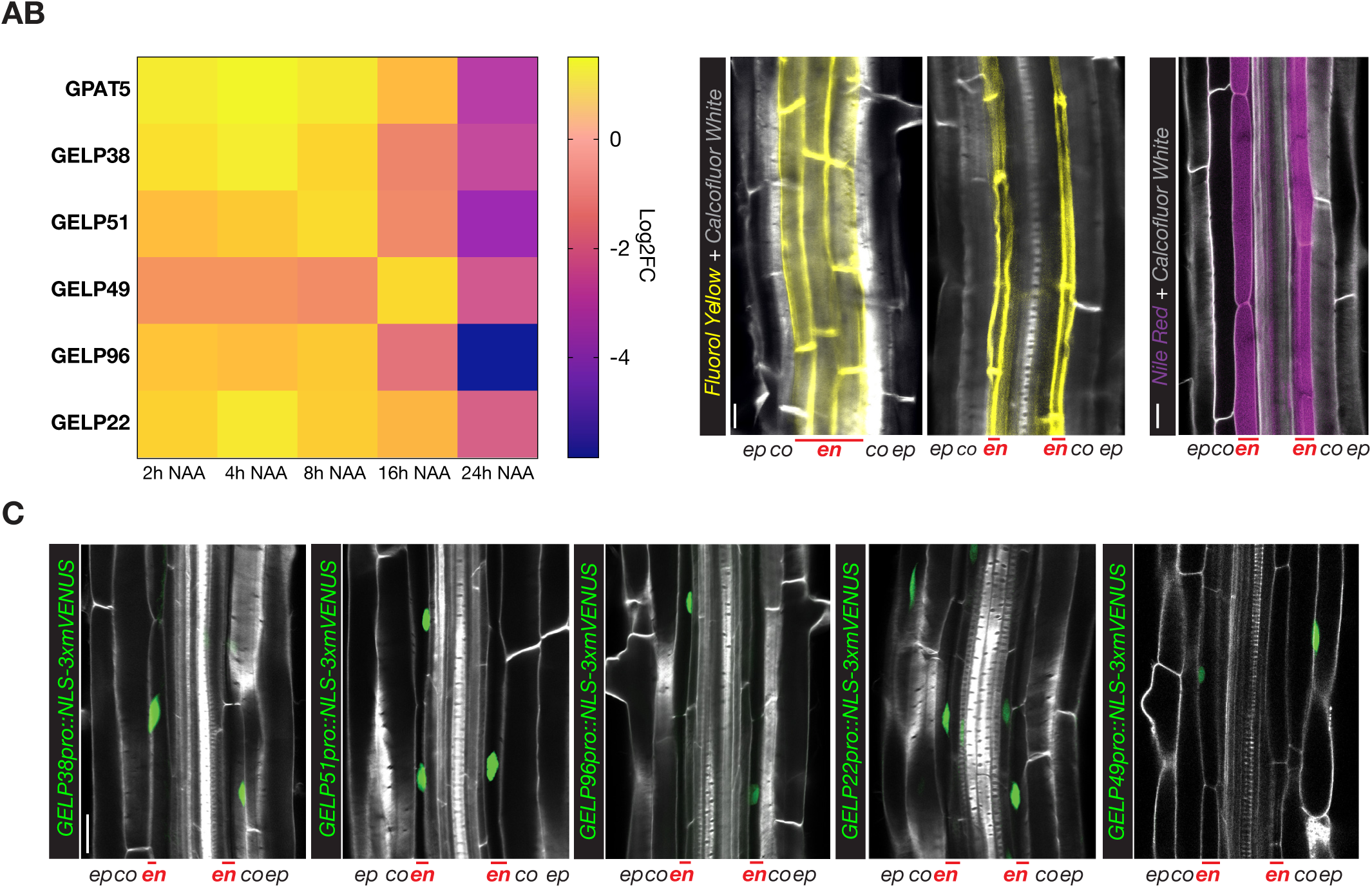
A cluster of five, auxin repressed, GELPs is expressed in the differentiated endodermis. **A**. Heatmap showing the differential expression of *GPAT5* and *GELP38, GELP51, GELP49, GELP96* and *GELP22* during the time course of NAA treatment (10 µM). **B**. Representative image of staining of suberin lamellae in the endodermis using Flurol Yellow (FY) (yellow) or Nile Red (magenta). **C**. Confocal images of root sections expressing transcriptional reporters for each of the GELPs mentioned in (**A**). NLS-3xmVENUS is shown in green, Calcofluor-White (CW) staining of cell walls in gray. Scale bar = 25 µm.

**Figure 4.**
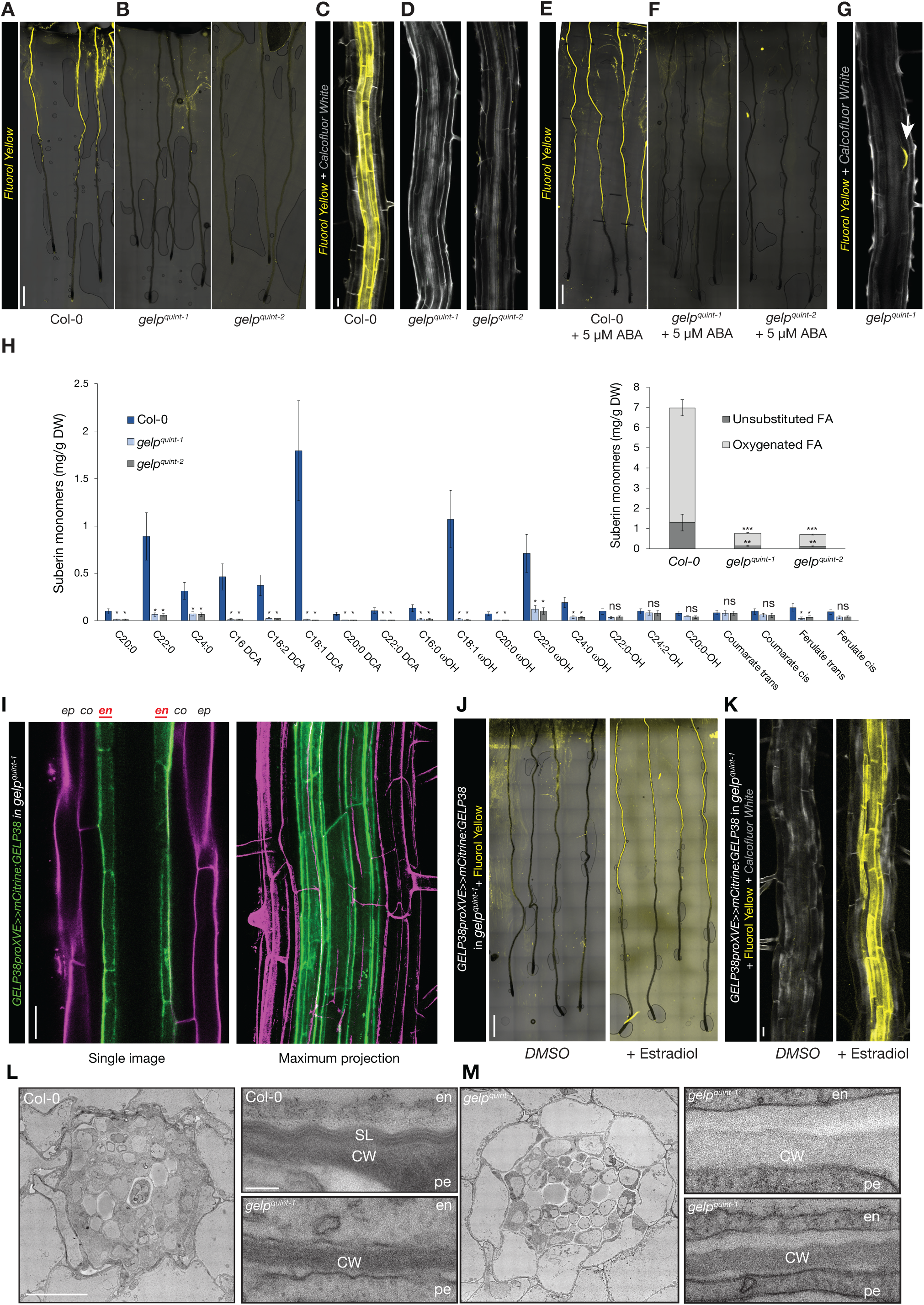
Suberin deposition strongly requires a cluster of auxin-repressed GELPs. **A**. Confocal images of Col-0 seedling roots stained with FY. **B**. Confocal images of *gelp*^*quint-1*^ and *gelp*^*quint-2*^ seedling roots stained with FY. Note absence of FY staining. **C** and **D**. Close up image of root sections of Col-0 (**C**) or *gelp*^*quint-1*^ and *gelp*^*quint-2*^ (**D**) stained by FY for suberin and CW for cell walls. **E** and **F**. ABA-induced increased suberin deposition in Col-0 roots (**E**) whereas (**F**) shows ABA cannot induce suberin deposition in roots of *gelp*^*quint-1*^ and *gelp*^*quint-2*^ mutants. **G**. FY staining showing that the cuticle layer protecting emerging lateral roots appears not to be affected in the *gelp*^*quint-1*^ mutant. **H**. Chemical analysis of the suberin content in roots of wild-type and *gelp*^*quint-1*^ and *gelp*^*quint-2*^ reveals a ∼85% decrease in total suberin monomers. Quantification of aliphatic and aromatic ester-bond suberin monomers isolated from 6-day-old roots of wild-type (Col-0) and *gelp*^*quint-1*^ and *gelp*^*quint-2*^ mutants. The graph shows the analysis of the principal suberin monomers and the inset shows the total monomers per genotype. Values represent the means ± SE, n = 4. Asterisks denote statistically significant differences to wild-type as determined by ANOVA and Tukey’s test analysis: ***p < 0.001; **p < 0.01; *p < 0.05, ns, not significant. **I**. Confocal image and maximum image projection of a root expressing *GELP38-XVEpro::mCITRINE:GELP38* (green) after β-Estradiol treatment (5 µM) counterstained with propidium iodide (magenta). **J**-**K**. Induction of *GELP38-XVEpro::mCITRINE:GELP38* restores suberin deposition in the roots of *gelp*^*quint-1*^. **L**-**M**. TEM micrographs of root cross sections showing presence of suberin lamellae in wild-type, and absence of suberin lamellae in cross sections of *gelp*^*quint-1*^ roots. Note that the structure of the endodermis of the *gelp*^*quint-1*^ mutant is much better preserved compared to wild-type. Scale bars in (**A, E, J**) = 500 µm, (**C, G, I, K**) = 25 µm. Scale bars in (**L**) and (**M**) for the whole root sections = 10 µm and for zoomed-in regions = 20 nm.

### Cell walls of endodermal cells in gelpquint-1 are devoid of suberin lamellae

TEM allows to directly observe the presence of suberin lamella in the endodermis. We therefore investigated the gelpquint-t mutant at the ultra-structural level. Whereas we could readily detect suberin lamellae deposition in endodermal cell walls of wild-type, we could not detect any indication of suberin lamellae formation in roots of *gelp*^*quint-1*^ (Figure 4L and M and Figure S6B). Instead, we observed a layer of low electron density in the cell wall of the gelpquint-1 mutant (Figure 4M and Figure S6B). This layer appears amorphous and bears no resemblance with the lamellar structures observed in wild-type. Hence, these results strongly support an absence of suberin in the endodermis of the *gelp*^*quint-1*^ mutant. Since it has been repeatedly demonstrated that roots with non-functional suberin barriers are more susceptible to elevated concentrations of salt, we also subjected 4 DAG seedlings to a mild salt stress (85 mM NaCl) for 10 days. Both *gelp*^*quint*^ mutants were much more affected by the salt stress compared to wild-type (Figure S6K-M). We observed much less emerged lateral roots and the fresh weight of the shoot was also significantly reduced (Figure S6K-M). These observations again strongly support the absence of a functional suberin barrier in the gelpquint mutants. Next, we checked whether the expression of known suberin biosynthesis-related genes was altered in the gelpquint-1 and gelpquint-2 mutants. Neither *GLYCEROL-3-PHOSPHATE SN-2-ACYLTRANSFERASE 5* (*GPAT5*), *HYDROLASE OF ROOT SUBERIZED TISSUE* (*HORST*), *ALIPHATIC SUBERIN FERULOYL-TRANSFERASE* (*ASFT*), *FAR1, FAR4* or *KCS2* were differentially expressed in the *gelp*^*quint-1*^ and *gelp*^*quint-2*^ mutants (Figure S6N). This excludes the possibility that the observed absence of suberin would be due to an indirect feedback regulation of suberin biosynthesis and supports a direct role for the five GELPs in suberin polymerization in the apoplast.

### A cluster of auxin-induced GELPs is required for suberin degradation

Besides a cluster of GELPs repressed by auxin treatment, we also identified a group of five GELPs (*GELP12, GELP55, GELP72, GELP73* and *GELP81*) that were induced by auxin (Figure 5A). Transcriptional reporters for GELP12, GELP55 and GELP72 revealed that their expression appeared to peak after 24 hr of NAA treatment, whereas reporters for *GELP73* and *GELP81* revealed a peak in expression after ∼4hr NAA treatment (Figure 5E and Figure S7D-F). In addition, both *GELP73* and *GELP81* were already expressed in the endodermis prior to lateral root formation (Figure 5E and Figure S7D).

**Figure 5.**
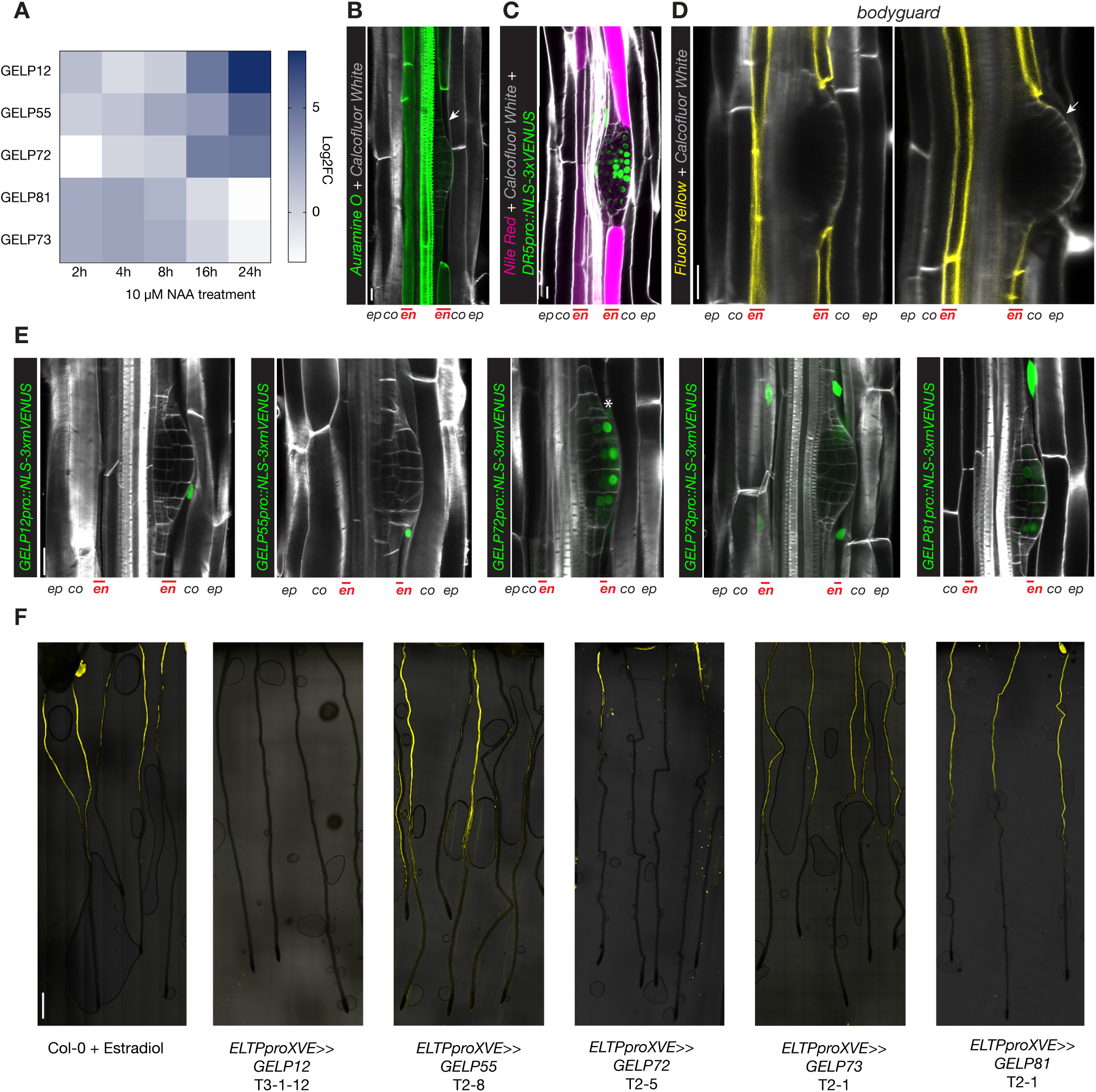
A cluster of auxin-induced GELPs is required for suberin degradation. **A**. Heatmap showing the differential expression of *GELP12, GELP55, GELP72, GELP81* and *GELP73* during the NAA (10 µM) time course. **B**. Use of Auramine O (green) to visualize degradation of suberin in the cell wall of endodermal cells overlying a lateral root primordium. **C**. Use of Nile Red staining (magenta) to visualize suberin degradation in the cell wall of endodermal cells overlying a lateral root primordium. Auxin signaling is visualized by *DR5pro::NLS-3xVENUS* (green). **D**. FY staining in roots of bodyguard, demonstrating normal presence of suberin lamellae, whereas the cuticle layer surrounding the lateral root primordium is discontinuous (arrow). **E**. Confocal images showing the expression patterns of the isolated auxin-upregulated GELPs during lateral root formation. NLS-3xmVENUS signal in green and Calcofluor White staining of cell walls is in gray. Asterisk indicates endodermal signal. **F**. FY staining on roots of Col-0 treated with ⊠Estradiol results in normal suberin pattern, whereas inducible endodermis-specific overexpression of *GELP12, GELP55* or *GELP72* results in degradation of suberin highlighted by absence of FY signal. The overexpression of *GELP73* and *GELP81* results in a normal suberin pattern similar to wild-type. Scale bars in (**B, C, D**) and (**E**) = 25 µm. Scale bar in (**F**) = 500 µm.

We demonstrated that the endodermal suberin gradually disappears and a layer of cutin is formed at the lateral root priordium (Figure 2), confirming previous observations (Berhin et al., 2019; Li et al., 2017). Thus, we hypothesized that, in contrast to the suberin biosynthesis GELPs that are auxin repressed, these five GELPs could instead regulate the removal of suberin in response to developmental and/ or environmental cues. First, we confirmed that also other fluorescent staining methods besides FY could visualize the local removal of suberin during lateral root formation. Both Auramine O and Nile Red can be used to visualize suberin in differentiated roots and both confirmed local degradation of suberin in cells overlying the lateral root (Figure 5B and C). Next, we assessed suberin staining in the *bodyguard* mutant. This α/β hydrolase was shown to be implicated in the establishment of the root cap cuticle (Berhin et al., 2019), but it was not clear whether it could also affect suberin accumulation in the endodermis. FY staining in bodyguard roots confirmed the effect on lateral root cap cuticle, but suberin degradation appears to be normal (Figure 5D and Figure S7B and C). When characterizing the expression patterns of the five putative suberin degrading GELPs, we observed three different expression patterns. *GELP12* and *GELP55* marker lines showed expression in the cortex or no signal in absence of lateral root formation, but were induced in endodermal cells overlying the lateral root from stage I to IV (Figure 5E and Figure S7D). *GELP72* and *GELP73* were only weakly induced in endodermal cells overlying the lateral root primordium, but from stage III and onwards they were induced in the outer layer of the growing primordium (Figure 5E and Figure S7E). Finally, *GELP81* was already present in the endodermis prior to lateral root initiation and was induced in the overlying endodermal cells from stage I to IV. In addition, from stage IV and onwards *GELP81* was induced in the outer layer of the primordium similarly to *GELP72* and *GELP73* (Figure 5E and Figure S7C). In order to characterize a possible role in suberin degradation, we first undertook a gain-of-function approach by generating plant lines in which we could inducibly overexpress each GELP in the endodermis and assess whether this causes suberin removal using FY staining. Indeed, inducible expression of *GELP12, GELP55* and *GELP72* in the endodermis caused a disappearance of FY staining, whereas no clear effect on FY staining was observed when inducing *GELP81* and *GELP73* (Figure 5F and Figure S8A). We then tested whether single loss-of-function mutants of each individual GELP affects lateral root formation. We performed the root bending assay in order to synchronously induce lateral roots and to quantify the progression of lateral root development at 18hr and 42hr after gravistimulation (Lucas et al., 2008; Péret et al., 2012). Among the three GELPs that can degrade suberin based on FY staining, only one (*gelp72*) showed a delayed lateral root development in both alleles and time points (Figure 6D-G). Surprisingly, the single mutants of the *GELP73* and *GELP81* genes - that did not affect FY staining when overexpressed in the endodermis - did display a delayed lateral root emergence (Figure 6H-K). Thus, a subset of auxin inducible GELPs can degrade suberin, possibly to facilitate lateral root emergence. Moreover, another subset of auxin inducible GELPs is also required for correct lateral root emergence, but may have functions unrelated to suberin degradation that we currently do not understand.

**Figure 6.**
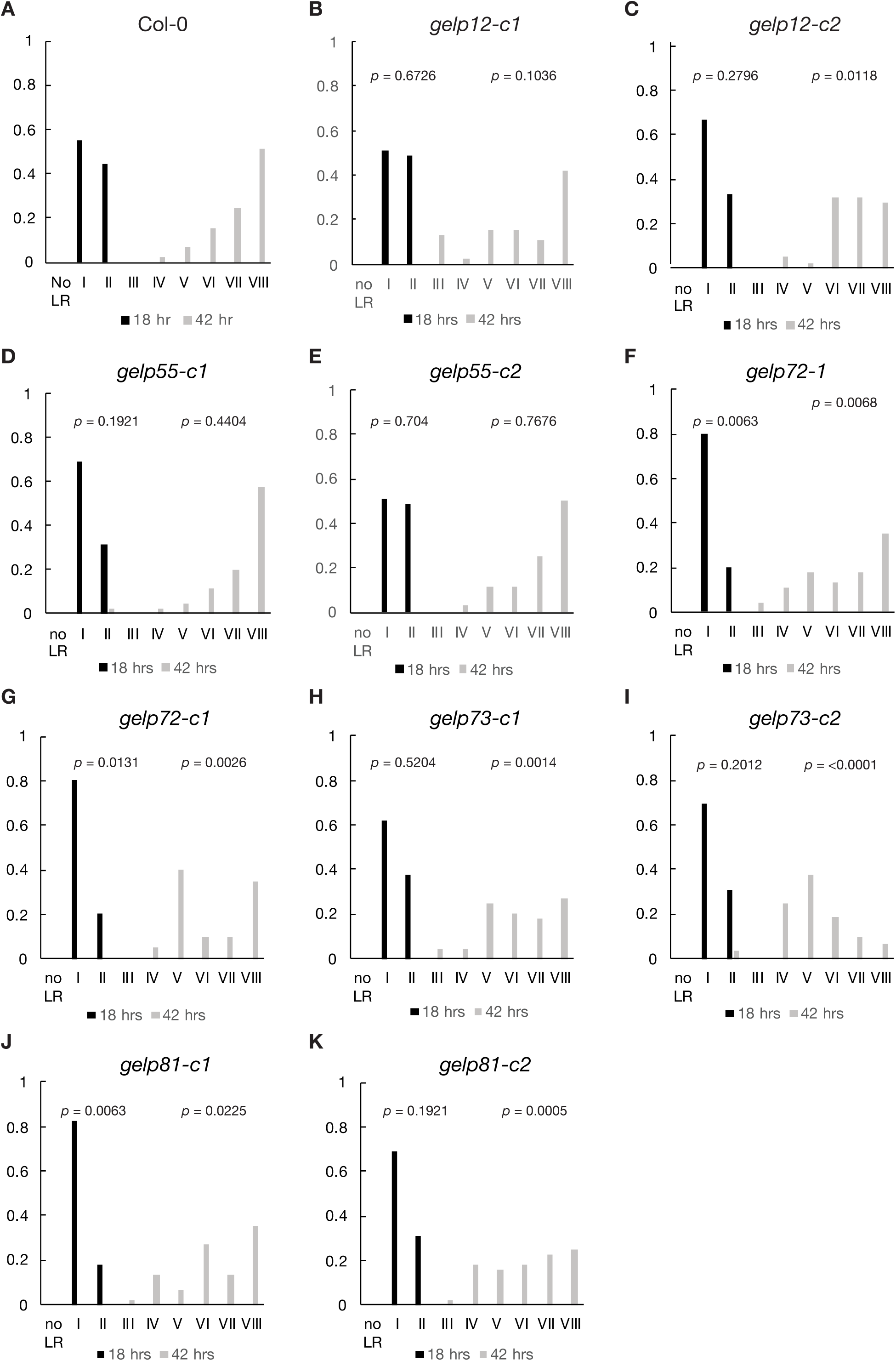
Auxin-inducible GELPs facilitate lateral root emergence. Gravistimulation-mediated induction of lateral root formation to functionally characterize the role of auxin-induced GELPs during lateral root formation. Staging of lateral root development was performed at 18hr and 42hr after gravistimulation. **A**. Col-0. **B** and **C**. *gelp12-c1* and *gelp12-c2*. **D** and **E**. *gelp55-c1* and *gelp55-c2*. **F** and **G**. *gelp72-1* and *gelp72-c1*. **H** and **I**. *gelp73-c1* and *gelp73-c2*. **J** and **K**. *p* values are indicated. A *p* value below 0.05 indicates a statistically significant difference as determined using the Pearsons’s χ^2^ test. Experiments were repeated three times with a minimal of 15 seedlings per genotype and time point.

## DISCUSSION

The auxin response of a differentiated tissue is very distinct from whole organ responses. We have shown in this study that differentiated root cells show response to auxin treatment. We achieved the strong enrichment for auxin responses in our specific cell type, the endodermis, by using different, tissue-specific auxin signaling mutants (Figure 1). A similar approach has also been employed to better understand auxin signaling during Arabidopsis embryogenesis (Schlereth et al., 2010). Comparing our data set to those of other auxin induced transcriptomes derived from whole roots or sections (Lewis et al., 2013; Voß et al., 2015) revealed that there was no clear overlap between these different datasets (Figure 2). This illustrates the importance of obtaining auxin-response profiles at cell type-specific resolution. It has been reported that nucleo-cytoplasmic partitioning of Auxin Response Factors was causing an attenuated auxin responsiveness in differentiated root tissues (Powers et al., 2019). While we cannot exclude an overall attenuation of auxin responses, we observed a similar induction of *SHY2pro::NLS-3xmVENUS* along the differentiated endodermis, suggesting a similar auxin responsiveness of the endodermis along the root, at least in the differentiated zone. Moreover, while qualitatively very different from other auxin response profiles including meristematic tissues, our endodermis-focused response profile of differentiated root sections revealed differential expression of close to a thousand genes at significant amplitudes, not supporting the view of a strong attenuation of auxin responses in differentiated tissues.

Moreover, our functional validation demonstrates that the endodermal auxin response profile obtained is reliable: 24 out of 27 genes tested either showed endodermal or lateral root-associated expression and among the GELP family members, their expression dynamics successfully predicted their respective roles in suberin biosynthesis or degradation.

### GDSL-domain containing genes are required for suberin homeostasis

Although the Arabidopsis genome contains more than 100 GDSL-domain containing genes, only a few of them have been functionally linked to cell wall modification or have been demonstrated to be able to synthesize or degrade cell wall polymers (Naseer et al., 2012; Philippe et al., 2020; Yeats et al., 2012). Several suberin biosynthetic genes as well as transcriptional regulators have been identified in recent years (Beisson et al., 2007; Cohen et al., 2020; Kosma et al., 2014; Yadav et al., 2014). Although suberin is chemically similar to cutin, there are significant differences in its composition and its deposition within the cell wall and we still do not know how suberin is polymerized and degraded. The finding that the GDSL-domain containing protein *CUTICLE DESTRUCTION FACTOR 1* (*CDEF1*) is able to degrade suberin when overexpressed in the endodermis already hinted at the capacity of these types of genes to regulate suberin homeostasis (Naseer et al., 2012). However, till now, no GDSL-domain containing genes responsible for the polymerization of suberin in the endodermis have been identified. Our study now indicates that this was due to a high degree of functional redundancy. Only when all five auxin-repressed GELPs were knocked-out did we obtain roots without detectable suberin (Figure 4). All five of the identified GELPs are predicted to be expressed in the endodermis and secreted, supporting a role in cell wall modification.

It was demonstrated that ABA treatment induces suberization in both the endodermis and cortex (Barberon et al., 2016) and the fact that all five GELPs required for suberin polymerization were induced by ABA treatment (Figure S4), further support their specific role in suberization. Although the chemical analysis revealed a ∼85% reduction in suberin monomers, it did not hint towards potential substrates for the biosynthetic GELPs. The ultra-structural studies on the roots of the *gelp*^*quint-1*^ mutant also did not show evident agglomerations of unpolymerized material. Rather, a distinct, homogenous zone of low electron density was observed in place of the suberin lamellae. Whether this zone is composed of the remnant, non-polymerized lipidic suberin components (possibly becoming extracted during TEM processing) or whether this is matrix material of unknown, potentially polysaccharide nature remains to be investigated. Nevertheless, our data clearly indicate that the five auxin-repressed GELPs are absolutely required for endodermal suberin formation. In addition, we could show, using a functional *GELP38:mCitrine* fusion, that the GELPs are localized and active in the apoplast of the endodermis. Due to the demonstrated in vitro activity of CD1, a homologous gene family member regulating cutin polymerization, we consider that the five GELPs we have identified are suberin synthases, mediating polymerization of suberin precursors in the cell wall.

In the same dataset from which we identified the GELPs responsible for suberin polymerization, we additionally identified five GELPs that displayed opposite behavior, being induced by auxin treatment. Our gain- and loss-of-function approaches show that the relationship between suberin degradation and lateral root emergence seems to be a rather complex one. Although we could demonstrate suberin degradation for three of the five auxin-induced GELPs, only one of these, *GELP72*, appeared to play a role in lateral root emergence on its own (Figure 6B-G). The idea that suberin needs to be degraded in order to facilitate lateral root emergence fits with the reduced number of lateral roots observed in the *abcg2*/*abcg6*/*abcg20* mutant that has higher levels of suberin in its roots (Yadav et al., 2014). It might well be that a defect in suberin degradation interferes with the spatial accommodation responses in the endodermis, resulting in a delayed emergence. Interestingly, both of the auxin-induced GELPs for which we could not show a direct role in suberin degradation (*GELP73* and *GELP81*) also displayed strong effects on lateral root emergence (Figure 6H-K). Although the phenotypes clearly suggest that they are involved in suberin homeostasis, follow-up studies are required to reveal if and how they modify suberin. Both genes appear to peak rather early during lateral root formation (Figure 5A) and could therefore also be involved in the formation of the cuticle at the lateral root primordium that protects the newly formed primordium. Indeed, defects in this structure also result in altered lateral root emergence (Berhin et al., 2019). We observed that several members of the group of auxin-inducible GELPs, besides being expressed in the endodermis, were also induced in the outer layer of the lateral root primordium (Figure 5). This would suggest that both the primordium and the endodermis contribute to suberin degradation and the concomitant formation of the lateral root cuticle. The fact that straight-forward overexpression of a single GELP can be sufficient to degrade suberin, fits to the idea that regulation of GELP expression is important to coordinate the degradation of the suberin lamellae and synthesis of the new cuticle structure. In this aspect it would be interesting to test if there is a functional hierarchy in the group of suberin degraders.

### GDSL-domain containing genes: an untapped potential for better understanding cell wall modifications and adaptation to different soil types

As mentioned before, the Arabidopsis family of GDSL-domain containing proteins contains more than hundred members, for which little functional information is available (Shi et al., 2011; Yeats et al., 2012). Here we demonstrate that five members belonging to this family are essential for suberin formation. Thus, it deserves more attention as its members are likely to perform multiple central biological roles in cell wall modifications during plant development or during adaptation to different stress conditions. Although the functional redundancy within this family is significant, it can nowadays be overcome by modern, gene-editing technologies, as we have shown with the cluster of GELPs required for suberin polymerization. We predict that the suberin mutant described in this work will be a valuable tool to better understand the process of suberin and cutin formation. Using endodermal specific promoters, we can now quickly test which additional GELPs can synthesize suberin. In addition, it is now straightforward to test if cutin synthases such as CD1 are specific for cutin or whether they can also synthesize suberin. The example of the suberin degrading subgroups moreover demonstrates that not all GELPs act redundantly. We are convinced that the GELPs in Arabidopsis and other plants provide a rich, but still untapped potential, to better understand and manipulate cell walls during plant development.

## Experimental Procedures

### Plant Material

For all experiments, Arabidopsis thaliana ecotype Columbia (Col-0; wild-type) was used. Gene numbers, mutants, and transgenic lines used and generated in this study are described in Supplemental Experimental Procedures. The primers used for genotyping and qPCR-based verification of T-DNA lines are indicated in Table S5.

### Plant growth conditions

For all experiments, plants were germinated on solid half-strength Murashige and Skoog (MS) medium without addition of sucrose. Seeds were surface sterilized, sown on plates, incubated for 2 days at 4°C for stratification, and grown vertically in growth chambers at 22°C, under continuous light (100 µE). The microscopic analyses (FDA uptake, FY staining, PI uptake, confocal microscopy) were performed on 5 or 7-day-old seedlings.

### Bending experiments

Seeds of wild-type Col-0 and gelp mutants were plated on half strength MS containing 120 × 120 x 17 mm square Petri dishes, stratified in the dark at 4°C for 2 days, and grown at 22 °C under constant light (100 µE). Lateral root stages were determined after plates with 4-day-old seedlings were rotated 90° degrees and grown for 18 hr and 42 hr for synchronized lateral root induction. After bending for 18hr and 42hr, the roots were cleared as described by (Voß et al., 2015) and mounted in 50% glycerol. Determination of lateral root stage in the bent region was done using an upright microscope with differential interference contrast optics. Experiments were repeated three times and each replicate had at least 15 seedlings.

### Generation of transgenic lines

*CASP1pro::shy2-2* was crossed into *slr-1* mutant to produce *CASP1::shy2-2*/*slr-1* line. For generating marker lines and overexpression constructs, the In-Fusion Advantage PCR Cloning Kit (Clontech) and Gateway Cloning Technology (Invitrogen) were used. All constructs were transformed by heat shock into Agrobacterium tumefaciens GV3101 strain and then transformed into plants by floral dipping (Clough and Bent, 1998). At least 10 independent transgenic lines were analyzed for expression patterns, and 1 line showing a representative signal and normal segregation was selected for further studies. For transcriptional reporters, the promoter regions were PCR-amplified from Col-0 genomic DNA and cloned into pDONRP4-P1R (www.thermofisher.com). The resulting plasmids were recombined together with pDONRL1-NLS-3xmVENUS-L2 (Gasperini et al., 2015) and the destination vector pFR7m24GW or pFG7m24GW, containing the FastRed or FastGreen cassettes for transgenic seed selection respectively, to create the final *PROMOTER::NLS-3xmVENUS* expression clones. To be able to induce expression of individual GELPs in the endodermis, the corresponding pDONR221_L1-GELP-L2 clones were created. The resulting clones were recombined with the estradiol-inducible pDONR_P4-ELTPproXVE-P1R (Andersen et al., 2018) and destination vector pB7m24GW, to produce *ELTPproXVE::GELP* overexpression lines. To generate *GELP38proXVE*>>*GELP38:mCitrine*, the promoter region of GELP38pro was PCR-amplified from Col-0 genomic DNA and cloned into linearized p1R4-ML:XVE (Siligato et al., 2016) with KpnI enzyme by Infusion (Takara) cloning to produce the inducible GELP38proXVE promoter clone. The resulting clone was recombined together with pDONR221_L1-GELP38-L2, pDONR_R2-mCitrine-L3 and pFG7m34GW to produce *GELP38proXVE*>>*GELP38:mCitrine* construct. See Table S3 and Table S4 for details about the primers used for cloning. The primers used for generating single and multiple CRISPR/Cas9 mutants are indicated in the Table S7. See Table S3 and Table S4 for details about the primers used for cloning. The primers used for generating single and multiple CRISPR/Cas9 mutants are indicated in the Table S7.

### Hormonal treatments

Abscisic acid (ABA) was stored as a 50 mM stock solution in methanol. When seedlings were subjected to short-term 10 µM ABA treatment, the transfer was done when the seedlings were 4-days-old. β-Estradiol was prepared as 100 mM stock in DMSO. In case of β-Estradiol treatment, the seedlings were directly germinated on the media containing 5 µM Estradiol. For salt experiments, the seedlings were grown on half-strength MS medium and transferred to 85mM NaCl for 10 days. In case of auxin (NAA) treatment for RNAseq experiments, the seedlings were first grown on half-strength MS medium and then transferred to 10 µM NAA for 4, 8, 16 and 24 hours. At each time point the, the shoots and the root tips were removed.

### Chemical analysis of suberin

Chemical analysis of suberin was performed on six-day old seedlings. Prior to analysis we confirmed that the used growth conditions did not affect the phenotype of *gelp*^*quint-1*^ and *gelp*^*quint-2*^ via FY staining. We used the protocol for the determination of ester-bond lipids as described by Berhin et al. (2019). In brief, 200 mg of seeds were grown on nylon mesh (200 mm pore size). After six days, the roots were shaved off after flash freezing and extracted in isopropanol/0.01% butylated hydroxytoluene (BHT). They were then delipidized three times (1 h, 16 h, 8 h) in each of the following solvents, i.e., chloroform-methanol (2:1), chloroform-methanol (1:1), methanol with 0.01% BHT, under agitation before being dried for 3 days under vacuum. Depolymerization was performed by base catalysis (Li-Beisson et al., 2013). Briefly, dried plant samples were transesterified in 2 mL of reaction medium. 20 mL reaction medium was composed of 3 mL methyl acetate, 5 mL of 25% sodium methoxide in dry methanol and 12 mL dry methanol. The equivalents of 5 mg of methyl heptadecanoate and 10 mg of ω-pentadeca-lactone/sample were added as internal standards. After incubation of the samples at 60°C for 2 h 3.5 mL dichloromethane, 0.7 mL glacial acetic acid and 1 mL 0.9% NaCl (w/v) Tris 100 mM pH 8.0 were added to each sample and subsequently vortexed for 20 s. After centrifugation (1500 g for 2 min), the organic phase was collected, washed with 2 mL of 0.9% NaCl, and dried over sodium sulfate. The organic phase was then recovered and concentrated under a stream of nitrogen. The resulting cutin monomer fraction was derivatized with BFTSA/pyridine (1:1) at 70°C for 1 h and injected out of hexane on a HP-5MS column (J&W Scientific) in a gas chromatograph coupled to a mass spectrometer and a flame ionization detector (Agilent 6890N GC Network systems). The temperature cycle of the oven was the following: 2 min at 50°C, increment of 20°C/min to 160°C, of 2°C/min to 250°C and 10°C/min to 310°C, held for 15 min. 3 independent experiments were performed with 4 replicates for each genotype, respectively, and a representative dataset is presented. The amounts of unsubstituted C16 and C18 fatty acids were not evaluated because of their omnipresence in the plant and in the environment.

### Fluorescence Microscopy

Confocal laser-scanning microscopy images were obtained using either a Zeiss LSM 880, Leica SP8 or Leica SP8-MP microscopes. For green and red fluorophores, the following excitation and detection windows were used: mVENUS/ GFP/FY/FDA 488 nm, 500-530 nm; mCITRINE 496 nm, 505-530 nm; PI 520 nm, 590-650 nm; Calcofluor White 405nm, 430-485 nm; Basic Fuchsin/Nile Red 561nm, 600-630nm. For multiphoton microscopy the following excitation and detection settings were used: mVENUS/GFP/FY/Calcofluor White 960nm, 435-485nm (Calcofluor White) and 500-550nm (mVENUS/GFP/FY). Methods for imaging the CS lignin and PI penetration were previously described (Fujita et al., 2020; Ursache et al., 2018). The details of modified methanol-based FY staining are presented in Supplemental Experimental Procedures. For visualization of FDA transport, chambered cover glasses (Thermo Scientific), were used where the roots were covered with a slice of agar and time lapses were made right after the application of FDA.

### Electron Microscopy

For chemical fixation, plants were fixed in glutaraldehyde solution (EMS, Hatfield, PA) 2.5% in phosphate buffer (PB 0.1 M [pH 7.4]) for 1h at RT and post fixed in a fresh mixture of osmium tetroxide 1% (EMS) with 1.5% of potassium ferrocyanide (Sigma, St. Louis, MO) in PB buffer for 1h at RT. The samples were then washed twice in distilled water and dehydrated in ethanol solution (Sigma, St Louis, MO, US) at graded concentrations (30%-40 min; 50% - 40 min; 70% - 40 min; 100% - 2×1h). This was followed by infiltration in Spurr resin (EMS, Hatfield, PA, US) at graded concentrations (Spurr 33% in ethanol- 4h; Spurr 66% in ethanol-4h; Spurr 100%-2×8h) and finally polymerized for 48h at 60°C in an oven. For the multiple mutant, ultrathin sections of 50 nm thick were cut transversally at 2 mm below the hypocotyl-root junction, using a Leica Ultracut (Leica Mikrosysteme GmbH, Vienna, Austria), picked up on a copper slot grid 2×1mm (EMS, Hatfield, PA, US) coated with a polystyrene film (Sigma, St Louis, MO, US). For lateral roots, ultrathin sections of 50 nm thick were cut longitudinally (transversally from main root). For High-Pressure freezing, plants were fixed in glutaraldehyde solution (EMS, Hatfield, PA) 2.5% in phosphate buffer (PB 0.1 M [pH 7.4]) for 1h at RT and post-fixed in a fresh mixture of osmium tetroxide 1% (EMS) with 1.5% of potassium ferrocyanide (Sigma, St. Louis, MO) in PB buffer for 1h at RT. The samples were then washed twice in distilled water before a High-Pressure Freezing step (HPF). For the High Pressure Freezing, 2mm long root pieces were cut below the hypocotyl junction region, and then placed in an aluminum planchet of 3mm in diameter with a cavity of 0.2mm (Art.241, Wohlwend GmbH, Sennwald, Switzerland) filled with Hexadecene (Merck KGaA, Darmstadt, Germany) covered with a tap planchet (Art.353, Wohlwend GmbH, Sennwald, Switzerland) and directly high-pressure frozen using a High-Pressure Freezing Machine HPF Compact 02 (Wohlwend GmbH, Sennwald, Switzerland). The samples were then dehydrated and infiltrated with resin at cold temperature using the Leica AFS2 freeze substitution machine (Leica Mikrosysteme GmbH, Vienna, Austria) with the following protocol: Dehydration in 100% Acetone (Sigma, St Louis, MO, US) at graded temperature (−90°C -10h; from -90°C to -60°C in 2h; -60°C for 8h; from -60°C to -30°C in 2h; -30°C -3h.) This was followed by infiltration in Spurr resin (EMS, Hatfield, PA, US) at graded concentration and temperature (30% -10h from -30°C to 0°C; 66% -10h from 0°C to 20°C; 100% -2X 10h at 20°C) and finally polymerized for 48h at 60°C in an oven. Ultrathin sections of 50 nm thick were cut transversally to the root, using a Leica Ultracut (Leica Mikrosysteme GmbH, Vienna, Austria), picked up on a copper slot grid 2×1mm (EMS, Hatfield, PA, US) coated with a polystyrene film (Sigma, St Louis, MO, US). Micrographs and panoramic were taken with a transmission electron microscope FEI CM100 (FEI, Eindhoven, The Netherlands) at an acceleration voltage of 80kV with a TVIPS TemCamF416 digital camera (TVIPS GmbH, Gauting, Germany) using the software EM-MENU 4.0 (TVIPS GmbH, Gauting, Germany). Panoramic were aligned with the software IMOD (Kremer et al., 1996).

### RNA-seq experiments

Seeds were surface sterilized, sown on plates, incubated 2 days at 4°C for stratification, and grown vertically in growth chambers at 22°C, under continuous light (100 µE) for 6 days. For each biological replicate (3 in total) 60 seedlings from each genotype were transferred to plates (20 seedlings per plate) containing ½ MS medium supplemented with 10 µM NAA and transferred back into the growth chamber. After the desired incubation period (2, 4, 8, 16 and 24hrs) seedlings were harvested after removal of the root apical meristem (∼3mm) and the shoot including hypocotyl and snap frozen in liquid nitrogen. RNA was extracted using a Trizol-based method. After RNase-free DNase (www.qiagen.com) treatment, RNA was cleaned-up using a RNeasy mini-elute kit (www.qiagen.com). RNA-seq libraries were prepared as described (Jan et al., 2019). In brief, RNA quality was assessed on a Fragment Analyzer (Advanced Analytical Technologies, Inc., Ankeny, IA, USA). RNA-seq libraries were prepared using 1000 ng of total RNA and the Illumina TruSeq Stranded mRNA reagents (Illumina; San Diego, California, USA) on a Sciclone liquid handling robot (PerkinElmer; Waltham, Massachusetts, USA) using a PerkinElmer-developed automated script. Cluster generation was performed with the resulting libraries using the Illumina TruSeq SR Cluster Kit v4 reagents and sequenced on the Illumina HiSeq 2500 using TruSeq SBS Kit v4 reagents. Sequencing data were processed using the Illumina Pipeline Software version 2.2.

### RNA-seq data processing and analysis

Data processing was performed by the Lausanne Genomic Technologies Facility using their in-house RNA-seq pipeline. Data analysis was done using an in-house RNA-seq pipeline that performed the following steps. Quality controls were applied for cleaning data for adapters and trimming of low-quality sequence ends. Cleaned data was aligned and read counts computed using two methods: STAR (Dobin et al., 2013) + HTSeq (Anders et al., 2015) and STAR + RSEM (Li and Dewey, 2011). First method generates gene counts and the second method generates isoform counts. TAIR10 genome and Ensembl 21 annotation were used. Additional quality controls were performed using R for inspecting the sample counts summary, pairwise sample correlations, clustering and sample PCA. Statistical analysis was performed for genes and isoforms with the Bioconductor package EdgeR (R version 3.4.0) for normalization and limma (R version 3.18.2) for differential expression. Two types of statistical tests were applied depending on the contrast model tested. A moderated t-test was used for each pairwise comparison in group t0 and group *slr-1* vs *CASP1pro::shy2-2/slr-1*. A moderated F-test was used for each time course model and their interaction. The result files contain one row per gene or transcript. Adjusted *p*-values have been computed for each comparison by the Benjamini-Hochberg method, controlling for false discovery rate (FDR). Genes were considered significant in further analysis if the adjusted *p*-value was equal or below 0.05 and the log2 fold change was ≥ 1. Further analysis has been conducted using R (v. 3.4.1). Heatmaps were generated using ComplexHeatmap (Gu et al., 2016) (v1.14.0) using pearson distance, and “average” for clustering. Non-supervised clustering of genes using kmeans (factoextra v1.0.4, https://github.com/kassambara/factoextra) suggested 3 clusters as optimal together. This represented a very low resolution and we looked into clusters of size 4-8 which contained slightly lower silhouette values. After testing manually multiple cluster suggestions we settled onto 7 inferred clusters based on the biologically most sensible separation. GO analysis was conducted using the package topGO [v. 2.28.0, weight01 algorithm; (Alexa and Rahnenfuhrer, 2019)]. GO annotations were obtained through org. At.tairGO (version 3.4.1). For the comparison with Lewis et al. (2013), the published series matrix file was obtained from the GEO archive and the differential gene-expression analysis repeated in order to obtain the expression of all genes. Results were compared to their published table of differentially expressed genes and found to be highly similar. A z-score based on the logFC value was calculated for both, our data-sets and the re-analyzed Lewis et al. (2013) data to make the different sets more comparable. For the comparison with Voß et al. (2015) we used directly the published table which included all expressed genes and calculated z-scores for the different time-points. We kept T0, T6, T9, T15 and T24 which resulted in 7145 differentially regulated genes with similar cut-off of *p*<0.05 and a fold-change of 2.

### qPCR analysis

For qPCR quantifications, the plants were grown on plates with half-strength MS medium covered with mesh. In case of quintuple gelp mutants, only root parts (around 100 mg) were collected and total RNA was extracted using a Trizol-adapted ReliaPrep RNA Tissue Miniprep Kit (Promega). For verifying the transcript level in single T-DNA lines, RNA extraction from whole seedlings was performed. Reverse transcription was carried out with PrimeScript RT Master Mix (Takara). All steps were done as indicated in manufacturer’s manual. The qPCR reaction was performed on an Applied Biosystems QuantStudio3 thermocycler using a MESA BLUE SYBR Green kit (Eurogentech). All transcripts were normalized to ADAPTOR PROTEIN-4 MU-ADAPTIN, AP4M (AT4G24550) expression. All primers used for qPCR are indicated in Table S6.

### HRM analysis of CRISPR mutants

HRM method was employed to screen for the mutants generated using CRISPR/Cas9-based method. Genomic DNA of selected Cas9-free, T2 generation plants, was extracted using CTAB DNA extraction method. The qPCR reaction was performed on Applied Biosystems QuantStudio3 thermocycler using a MeltDoctor HRM Master Mix, according to manufacturer’s indications (Applied Biosystems). HPLC-purified primers were used to generate an amplicon of around 200 base pairs. The results were analyzed using High Resolution Melt Software v3.1 (Thermo Fisher Scientific). The selected candidates were verified by sequencing. Primers used for amplification and sequencing the potential mutation sites are indicated in Table S8 and table S9.

### Quantification and Statistical Analysis

For quantifying the FY occupancy, confocal images were analyzed with the Fiji package (http://fiji.sc/Fiji) (Schindelin et al., 2012). Contrast and brightness were adjusted in the same manner for all images. The suberized regions of the roots were measured together with total root lengths to determine the percentage of suberin occupancy. All statistical analyses were done with the GraphPad Prism 8.0 software (https://www.graphpad.com/) or using the R package (http://www.r-project.org). One-way ANOVA was performed, and Tukey’s test was subsequently used as a multiple comparison procedure. For the analysis of lateral root development using the bending assay, we used a Pearsons’s χ^2^ test. Details about the statistical approaches used can be found in the figure legends. The data are presented as mean ± SD, and “n” represents number of plant roots. Each experiment was repeated at least 3 times.

## ACKNOWLEDGEMENTS

We would to thank Christopher Grefen and Marie Barberon for insightful discussion about GDSL-domain containing proteins, experimental approaches and feedback on the manuscript. We thank the Electron Microscopy Facility and Imaging Facility of the University of Lausanne and the Center of Microcopy and Image Analysis of the University of Zurich for excellent service and support. Work in the Geldner lab was supported by an ERC Consolidator Grant (GA-N: 616228-ENDOFUN), two consecutive Swiss National Science Foundation (Schweizerischer Nationalfonds zur Förderung der Wissenschaftlichen Forschung) grants (CRSII3_136278 and 31003A-156261). Robertas Ursache was supported by an EMBO Long-Term Fellowship (EMBO ALTF 1046-2015). Work in the Nawrath lab was supported by SNSF grant (310030_188672/1). Work in the Vermeer lab was supported by SNSF grants PP00P3_157524 and 316030_164086) and support from the University of Neuchâtel.

## AUTHOR CONTRIBUTIONS

R.U., N.G. and J.E.M.V. conceived, designed and coordinated the project. R.U., C.D.J.T.V, V.D.T., D.d.B., K.G., V.S., T.G.A. and J.E.M.V. performed all eperimental work. E.S.S., S.C. and S.P. analysed NGS data. J.E.M.V. wrote the first draft of the manuscript. R.U. T.G.A. C.N. N.G. and J.E.M.V. revised the manuscript and all authors were involved in the discussion of the work.

---

**Supplemental Figure S1.**
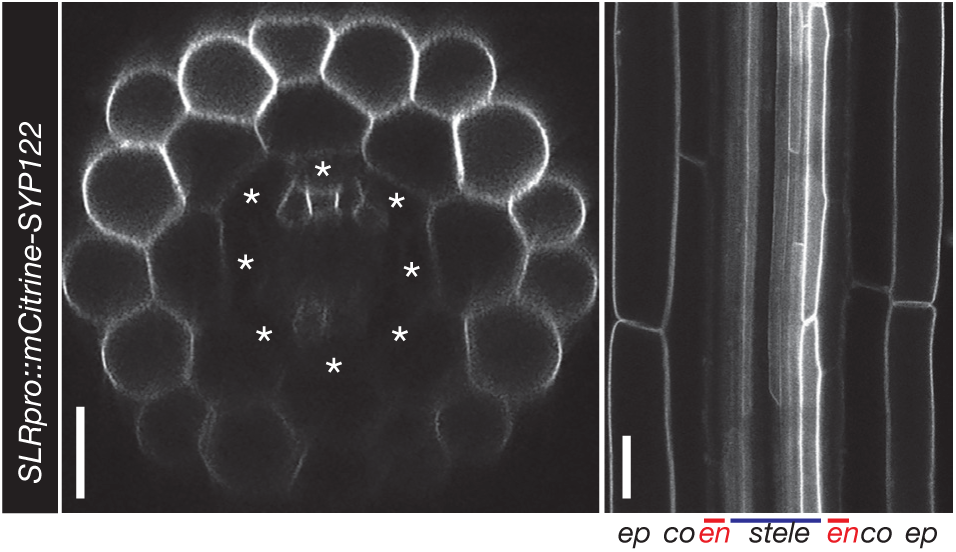
*SOLITARY ROOT* is not expressed in the endodermis. Confocal images of a root expressing *SLRpro::CITRINE:SYP122* confirming that SLR is expressed in the epidermis, cortex, pericycle and weakly in the stele, but not in the endodermis (indicated by asterisks). Scale bar = 25 µm.

**Supplemental Figure S2.**
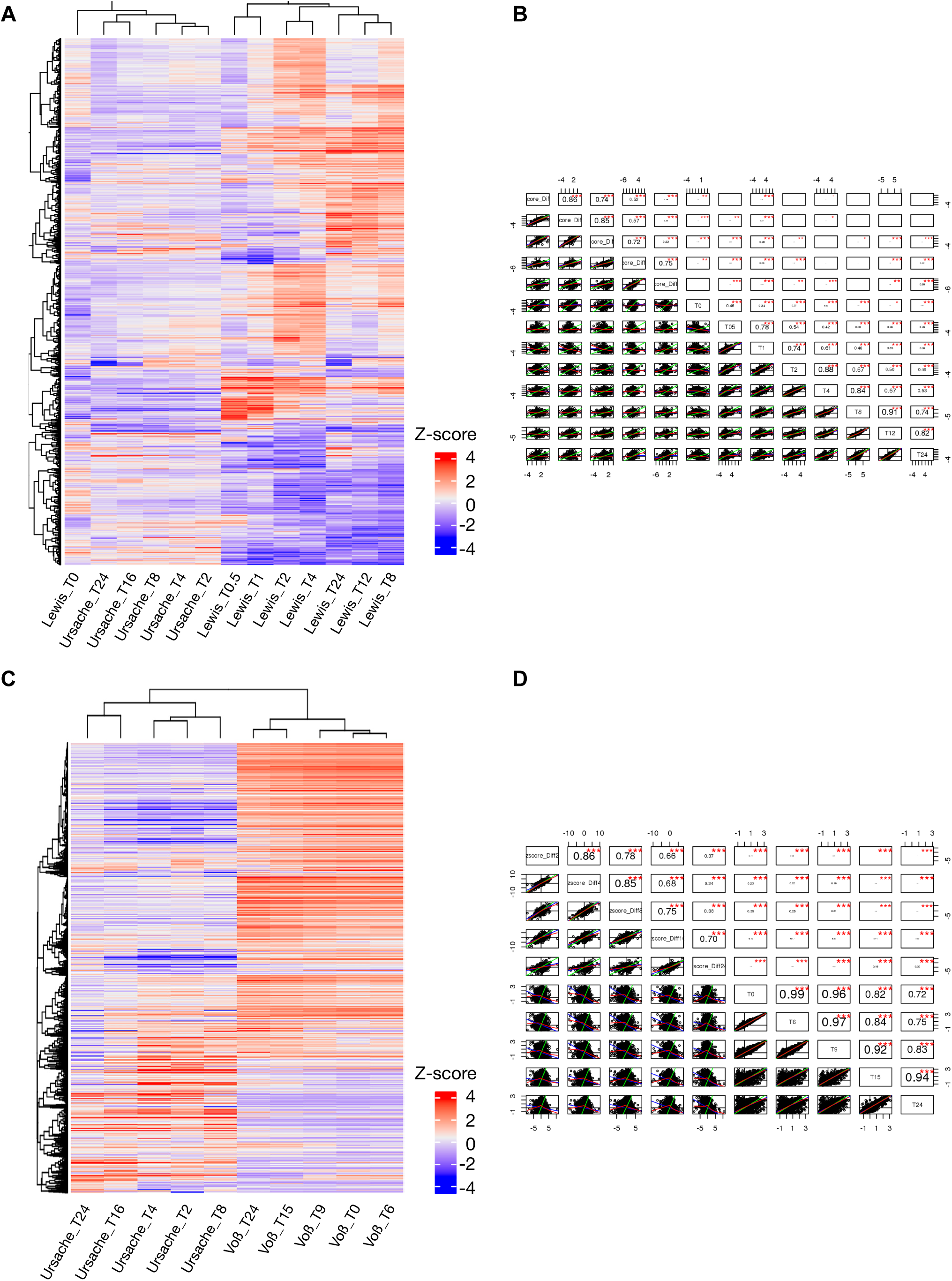
Differentiated endodermal cells have a distinct transcriptional response to auxin. **A** and **B**. Comparison of the current dataset (Ursache) with the data set of Lewis et al., (2013). **A**. Heatmap showing that both datasets cluster separately and do not have significant overlap, which is confirmed by the analysis of correlation between the two data sets (**B**). **C** and **D**. Comparison of the current dataset (Ursache) with the data set of Voß et al., (2015). C. Heatmap showing that both datasets cluster separately and do not have significant overlap, which is confirmed by the analysis of correlation between the two data sets (D).

**Supplemental Figure S3.**
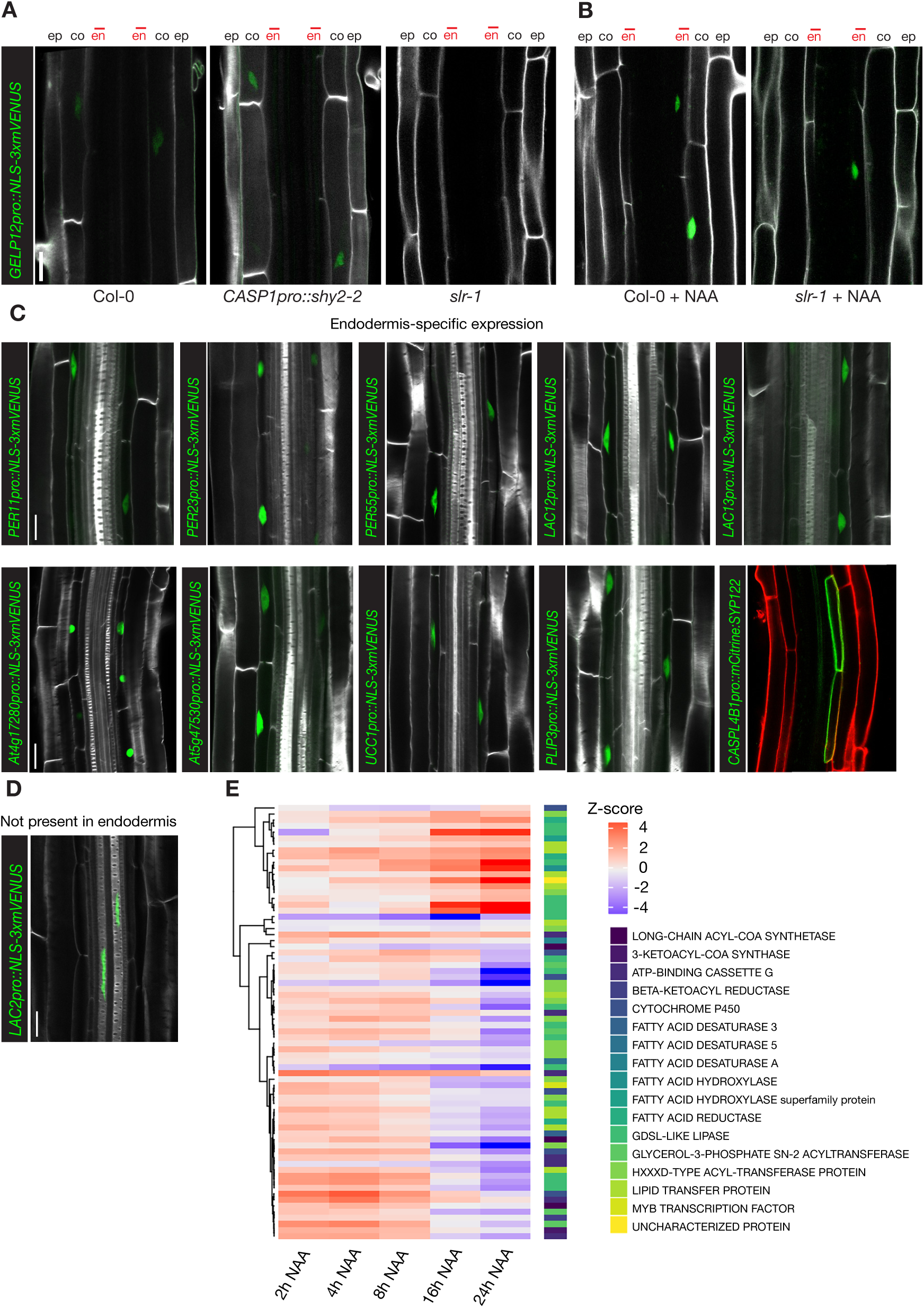
A high number of differentially regulated genes are expressed in the endodermis. **A**. Confocal images of *GELP12pro::NLS-3xmVENUS* expression in different genetic backgrounds, showing repression in *slr-1* under control conditions. **B**. Auxin treatment (10 µM NAA, 16hrs) results in induction of *GELP12pro::NLS-3xmVENUS* expression in the endodermis of *slr-1* roots. NLS-3xmVENUS signal is shown in green and CFW staining of cell walls is shown in grey. **C**. Confocal images of roots expressing transcriptional markers of candidate genes differentially expressed between *slr-1* and *CASP1pro::shy2-2*/*slr-1* roots and showing specific expression in the endodermis. **D**. Confocal images showing xylem-specific expression of *LAC2pro::NLS-3xmVENUS*. **E**. Heatmap showing the expression dynamics of suberin-related genes significant differentially expressed. NLS-3xmVENUS signal is shown in green, CFW staining of cell walls in gray and cell wall staining by PI in red. Scale bars = 20 µm.

**Supplemental Figure S4.**
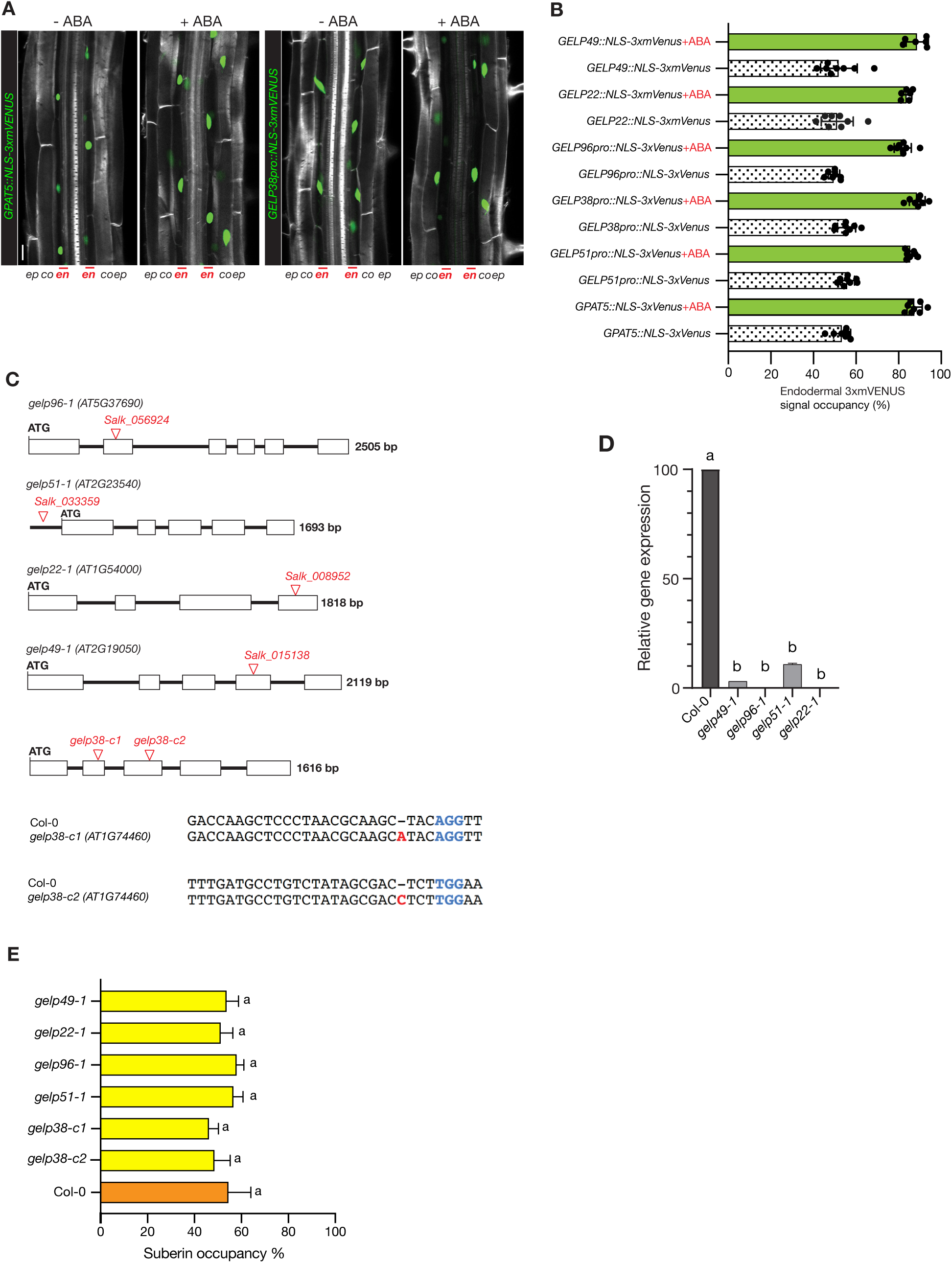
Expression of suberin biosynthesis-related GELPs is induced by ABA treatment. **A**. Confocal images showing effect of ABA treatment (1 µM, 24hr) on the expression domain of *GPAT5pro::NLS-3xmVENUS* and *GELP38pro::NLS-3xmVENUS* in Arabidopsis roots. **B**. Quantification of the effect of ABA treatment (1 µM, 24hr) on the expression of suberin biosynthesis-related GELPs identified as being repressed by auxin treatment. **C**. Schematic representation of the different single mutants of the suberin biosynthesis-related GELPs used in this study. **D**. qPCR results showing the effect of the T-DNA insertions of the suberin synthesis-related GELPs used in this study. Experiment was performed on three biological replicates. E. Quantification of suberin occupancy in the endodermis of the single mutants of the suberin biosynthesis-related GELPs using FY staining. Different letters in (**D**) and (**E**) (*p* < 0.05) indicate statistically significant differences between means by ANOVA and Tukey’s test analysis. ns, not significant. Scale bar in (**A**) = 25 µm.

**Supplemental Figure S5.**
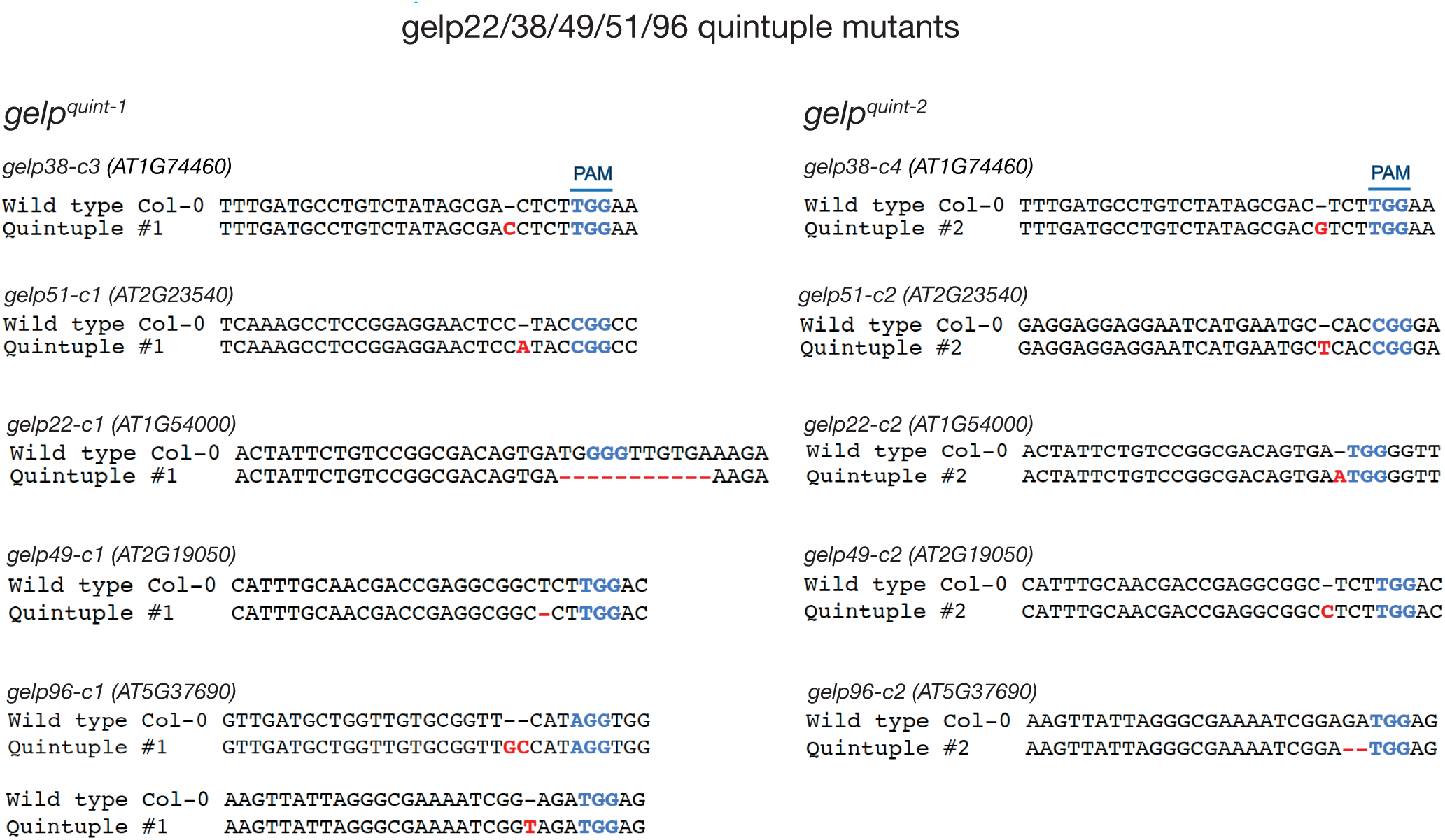
CRISPR/Cas9-mediated gelpquint mutants. Schematic representation of the mutations in the *gelp*^*quint-1*^ and *gelp*^*quint-2*^ mutants. The mutations are indicated in red and the PAM sites in blue.

**Supplemental Figure S6.**
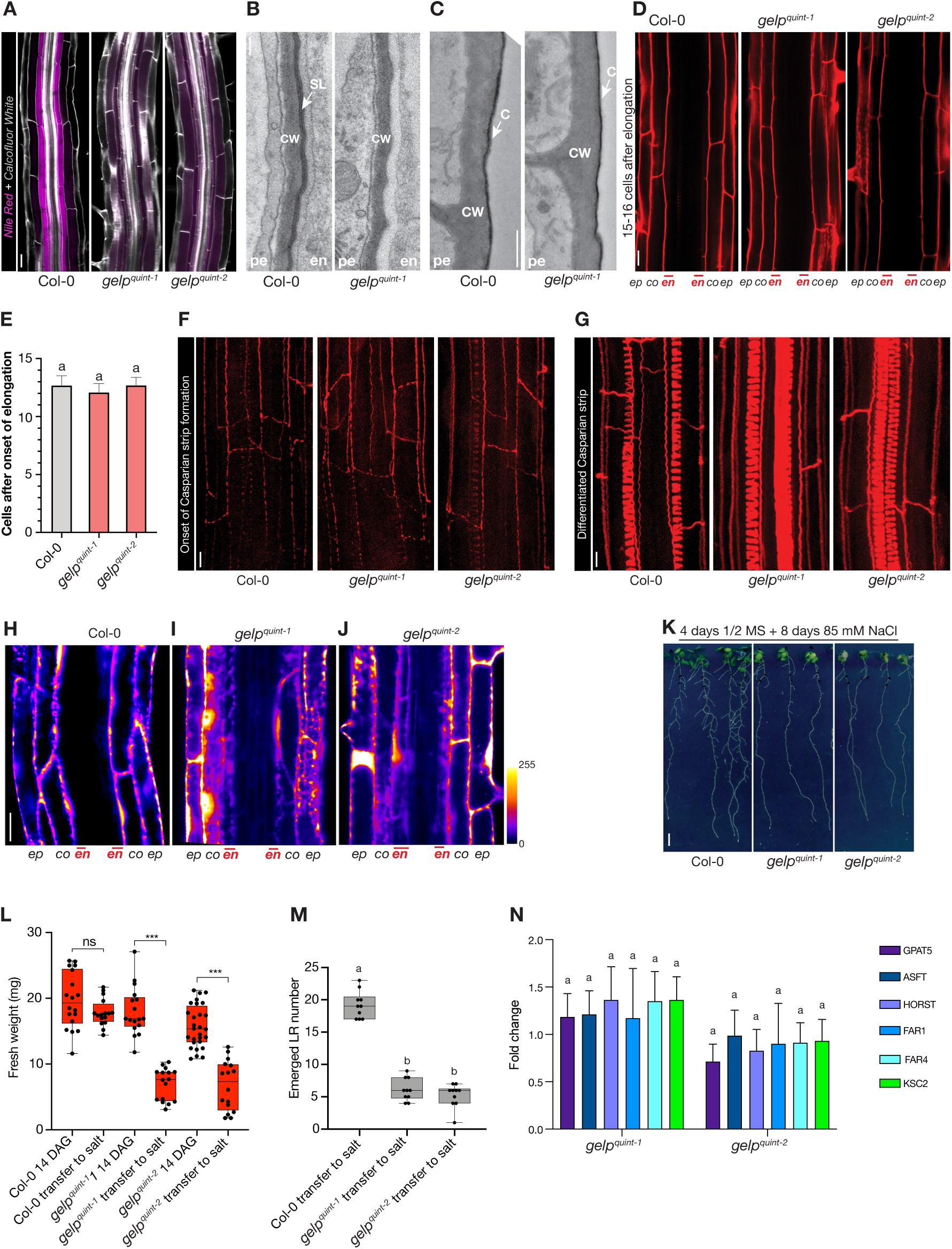
Extended characterization of the phenotype of *gelp*^*quint-1*^ and *gelp*^*quint-2*^ mutants. **A**. Nile Red staining of wild-type and *gelp*^*quint-1*^ and *gelp*^*quint-2*^ roots confirms the absence of suberin in the mutants. **B**. TEM micrographs of high-pressure frozen roots of wild-type and *gelp*^*quint-1*^ showing the absence of suberin lamella in the suberin biosynthesis mutant. **C**. TEM micrograph of high-pressure frozen roots of wild-type and *gelp*^*quint-1*^ showing that the lateral root cap cuticle is not affected in the suberin biosynthesis mutant. **D**. PI-mediated barrier assay on wild-type and *gelp*^*quint-1*^ and *gelp*^*quint-2*^ roots. **E**. Quantification of PI uptake in roots of wild-type and *gelp*^*quint-1*^ and *gelp*^*quint-2*^ seedlings. **F-G**. Basic Fuchsin staining of the Casparian strip in early and differentiated endodermal cells of wild-type and *gelp*^*quint-1*^ and *gelp*^*quint-2*^ roots. **H**. Fluorescein di-acetate (FDA) uptake assay in wild-type roots showing a suberin mediated block of uptake at the level of the endodermis. **I**. FDA uptake in assay in *gelp*^*quint-1*^ and (**J**) *gelp*^*quint-2*^ mutant roots, showing FDA uptake is not blocked at the level of the endodermis. **K**. Salt stress assay showing that *gelp*^*quint-1*^ and *gelp*^*quint-2*^ mutant seedlings are more sensitive to mild salt stress (85 mM NaCl) compared to wild-type. **L**. Quantification of the effect of prolonged salt stress on the fresh weight of wild-type and *gelp*^*quint-1*^ and *gelp*^*quint-2*^ seedlings. **M**. Quantification of emerged lateral roots in wild-type and *gelp*^*quint-1*^ and *gelp*^*quint-2*^ mutants after 12 days of exposure to salt. **N**. Quantification of the expression of known suberin biosynthetic genes in *gelp*^*quint-1*^ and *gelp*^*quint-2*^ mutants. Results are presented as fold-change compared to their expression levels in wild-type. Results were obtained from three biological replicates. Different letters in (**E**) and (**M**) (p < 0.001) and asterisks in (**L**) (*p* < 0.001) indicate statistically significant differences between means by ANOVA and Tukey’s test analysis. ns, not significant. Scale bars for (**A**), (**D**), (**F**), (**G**), (**H-J**) = 25 µm. Scale bars for (**B**) and (**C**) = 1 µm, for (**K**) = 5mm.

**Supplemental Figure S7.**
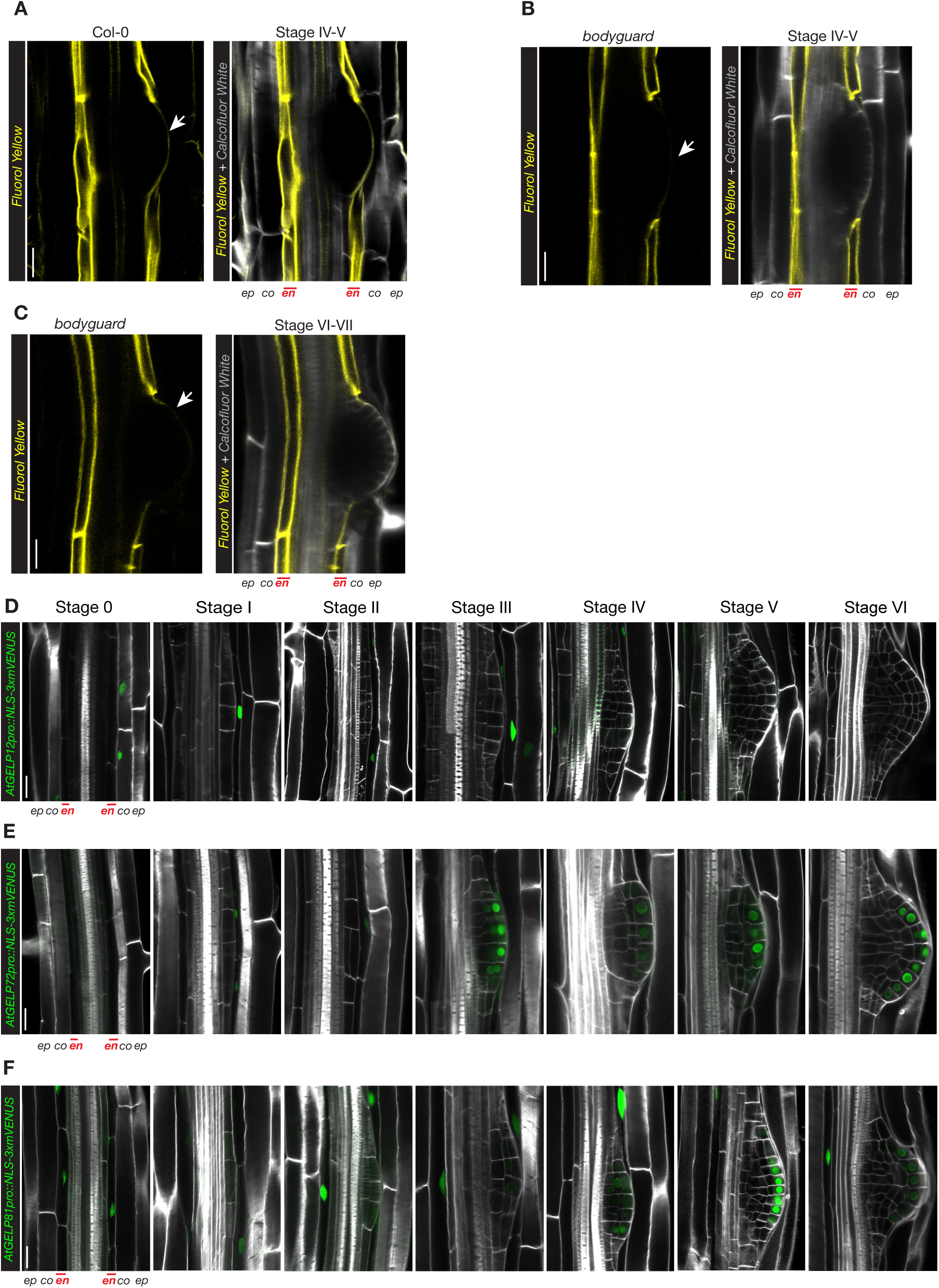
Auxin-upregulated GELPs show three distinct expression patterns. **A**. FY staining of wild-type Col-0 root at the site of lateral root emergence highlighting the presence of cuticle (indicated by arrow). **B**-**C**. FY staining of bodyguard mutant root at the site lateral root emergence. The absence of cuticle highlights the gap in FY staining in endodermis. **D**-**F**. Confocal images showing the expression patterns of *GELP12* (**D**), *GELP72* (**E**) and *GELP81* (**F**) during lateral root emergence. NLS-3xmVENUS signal is in green and Calcofluor White staining of cell walls is in gray. Scale bars = 25 µm.

**Supplemental Figure S8.**
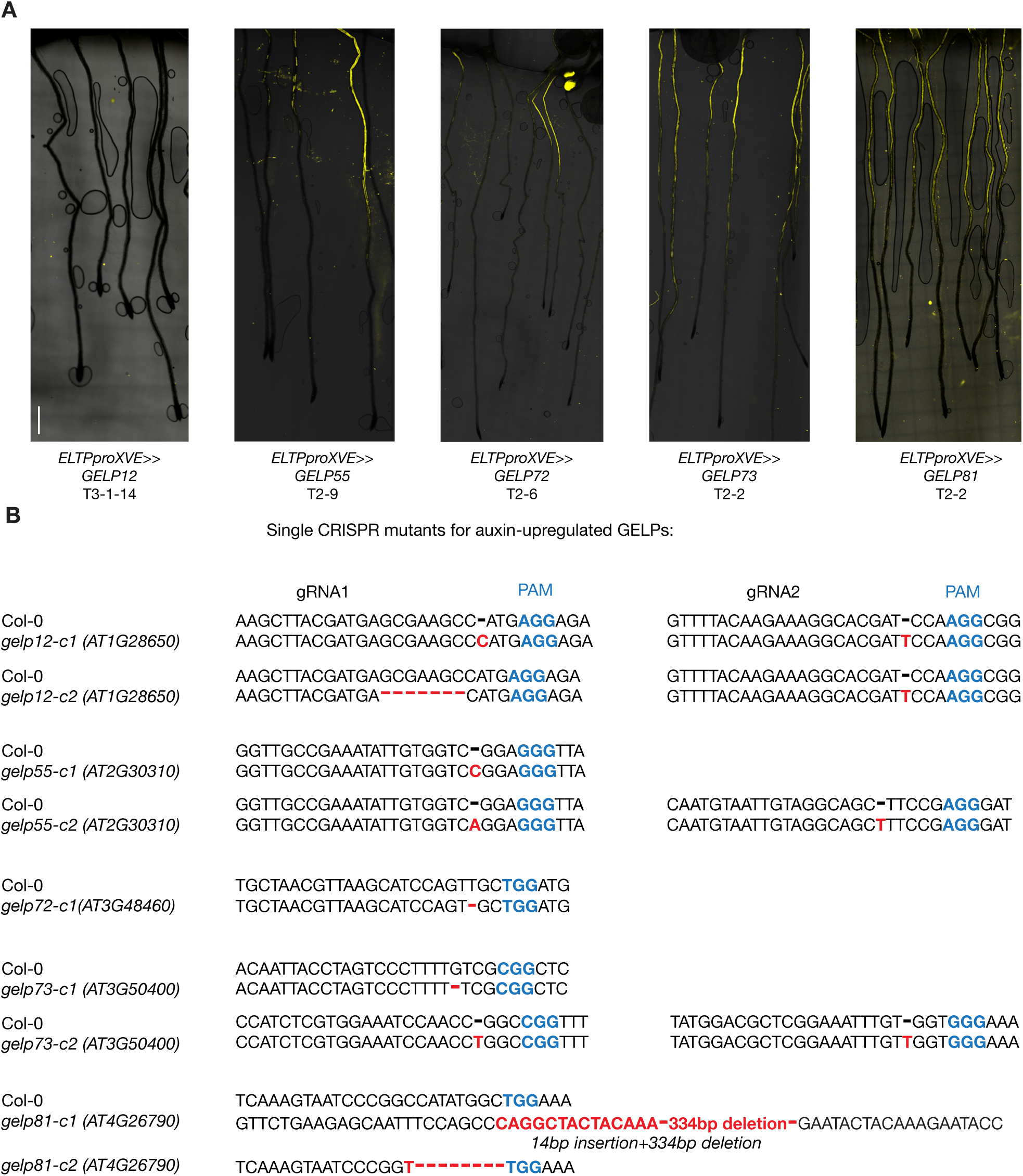
Overexpression of three auxin-induced GELPs leads to suberin degradation. **A**. FY staining on roots of Col-0 treated with β-Estradiol results in normal suberin pattern, whereas inducible endodermis-specific overexpression of *GELP12, GELP55* or *GELP72* results in degradation of suberin highlighted by absence of FY signal. The overexpression of *GELP73* and *GELP81* results in a normal suberin pattern similar to wild-type. **B**. Schematic representation of the mutations in the auxin-upregulated single GELP mutants. The mutations are indicated in red and the PAM sites in blue. Scale bars in (A) = 500 µm.

## Supplemental Experimental Procedures

### Plant material and constructions

The following T-DNA tagged and transgenic lines were used in this study: *gelp49* (SALK_015138C); *gelp51* (SALK_033359C); *gelp96* (SALK056924C); *gelp96* (SALK056924C) were requested from NASC center; *SHY2pro::NLS-3xmVENUS, CASP1pro::shy2-2, DR5::NLS-3xVENUS* (Vermeer et al., 2014); *GPAT5pro::NLS-3xmVENUS* (Ursache et al., 2018), *slr-1* (Fukaki et al., 2002) were described previously. The corresponding gene numbers are as follow: SLR, At4g14550; *SHY2*, At1g04240; *ELTP*, At2g48140; *GPAT5*, At3g11430; *SYP122*, At3g52400; *BDG*, At1g64670; *HORST*, At5g58860; *ASFT*, At5g41040; *FAR1*, At5g22500; *FAR4*, At3g44540; *KCS2*, At1g04220; *GELP12*, At1g28650; *GELP55*, At2g30310; *GELP72*, At3g48460; *GELP81*, At4g26790; *GELP73*, At3g50400; *GELP49*, At2g19050; *GELP51*, At2g23540; *GELP96*, At5g37690; *GELP38*, At1g74460; *GELP22*, At1g54000; *GELP103*, At5g45960; *PER10*, At1g49570; *PER11*, At1g68850; *PER23*, At2g38390; *PER28*, At3g03670; *PER55*, At5g14130, *PER59*, At5g19890; *LAC2*, At2g29130; *LAC12*, At5g05390; *LAC13*, At5g07130; *LAC16*, At5g58910; *UCC1*, At2g32300; *PLIP3*, At3g62590; *CASPL4B1*, At2g38480; *CASPL1A1*, At1g14160; *CYTOCHROME b561* and *DOMON DOMAIN-CONTAINING PROTEIN*, At4g17280; CYTOCHROME b561 and DOMON DOMAIN-CONTAINING PROTEIN, At5g47530.

### Methanol-based Fluorol Yellow staining of suberin in combination with Calcofluor White

For most experiments suberin lamellae were observed in 5 or 7-day-old roots using Fluorol Yellow (FY 088, SANTA CRUZ BIOTECHNOLOGY) staining. Seedlings were incubated in methanol at room temperature for at least three days, stained with FY 088 (0.01%, methanol) for 1 hour at room temperature, rinsed in methanol and counterstained with aniline blue (0.5%, methanol) at room temperature for 1 hour in darkness, washed, and visualized using 1-well chambered cover glass (ThermoFisher Scientific, Catalog Nr. 155361). In order to combine with Calcofluor White for cell wall staining, the seedlings were incubated first in Calcofluor White solution (0.1%, in methanol), for three days and stained with FY as described above.

---

**Supplemental Table S1.** Significant differentially expressed genes in *slr-1* versus *CASP1pro::shy2-2/slr-1* roots.

| Group | Time point | F-test Post Hoc classification |  |  |  |  |  |
| --- | --- | --- | --- | --- | --- | --- | --- |
|  |  | FDR < 0.05 |  |  | FDR < 0.05, Fold change > 2 |  |  |
|  |  | ALL | UP | DOWN | ALL | UP | DOWN |
| *slr-1* vs<br>*CASP1pro::shy2-2/slr-1*<br>timeseries<br>(10µM NAA) | 2 hr | 3043 | 1530 | 1513 | 865 | 469 | 396 |
|  | 4 hr | 2974 | 1523 | 1451 | 933 | 522 | 411 |
|  | 8 hr | 2736 | 1458 | 1278 | 824 | 478 | 346 |
|  | 16 hr | 2278 | 1267 | 1011 | 818 | 509 | 309 |
|  | 24 hr | 3093 | 1450 | 1643 | 1014 | 559 | 455 |

**Supplemental Table S2.**
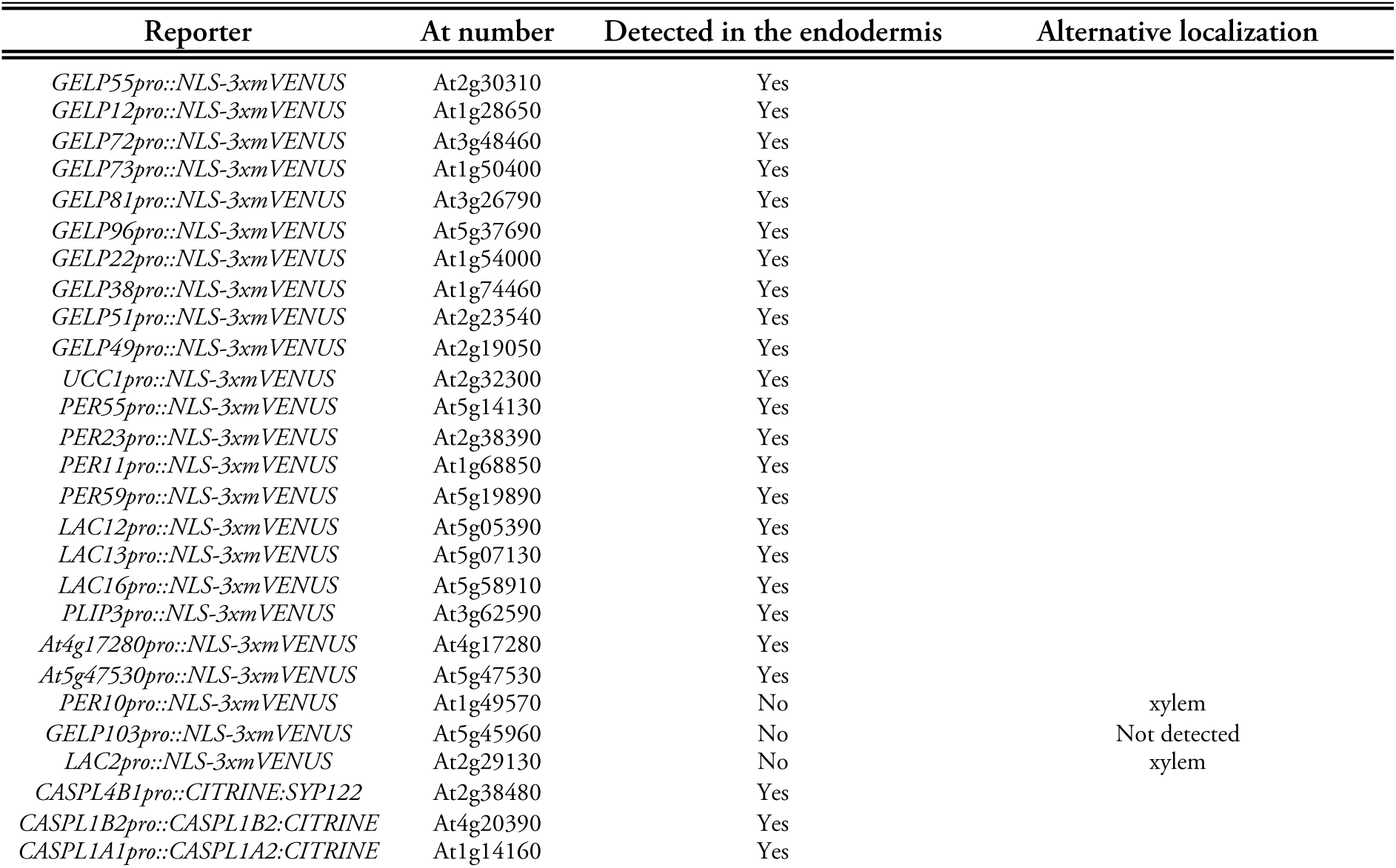
Transcriptional reporters used for confirmation of RNAseq data.

**Table S3.**
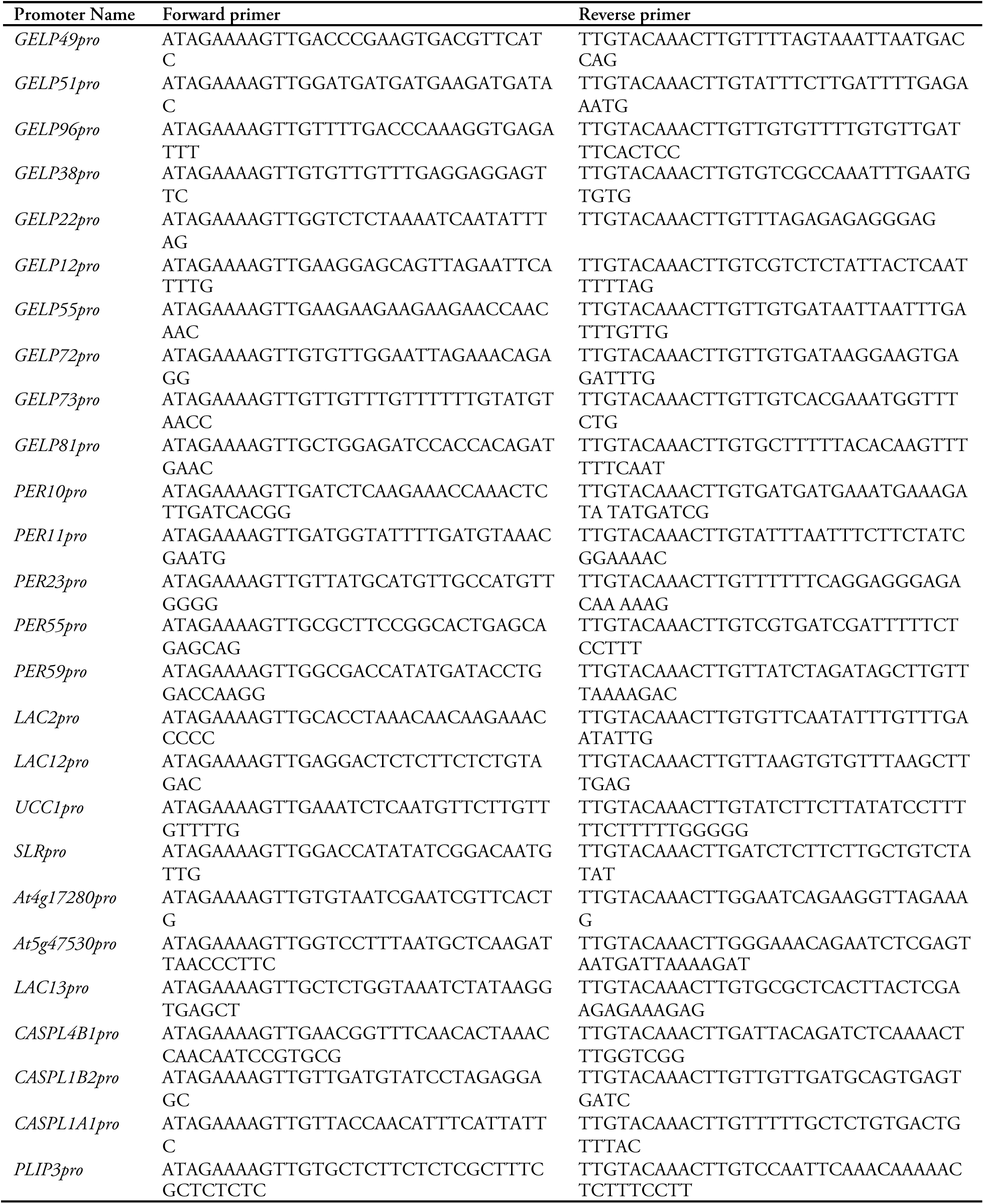
List of primers for cloning promoters into pDONR_P4R1 entry clone for Gateway assembly.

**Table S4.**
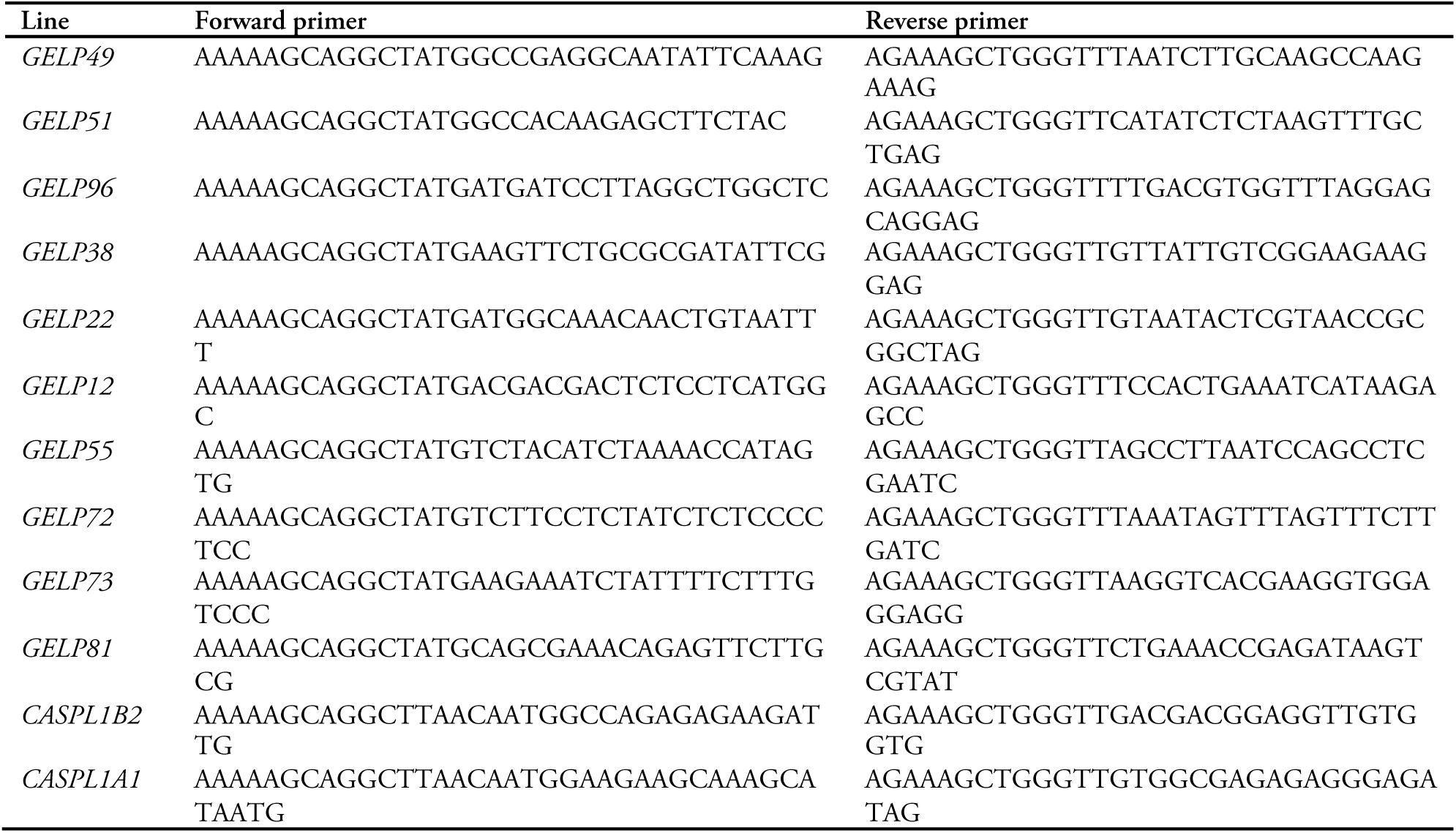
List of primers for cloning genomic fragments into pDONR221 Gateway entry clone.

**Table S5.**
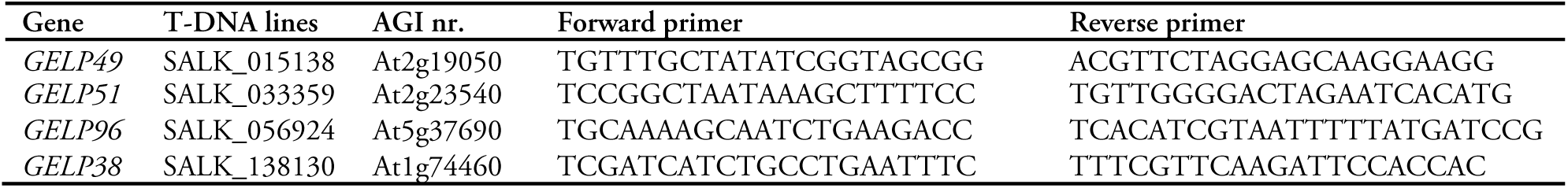
Primers for genotyping T-DNA lines.

**Table S6.**
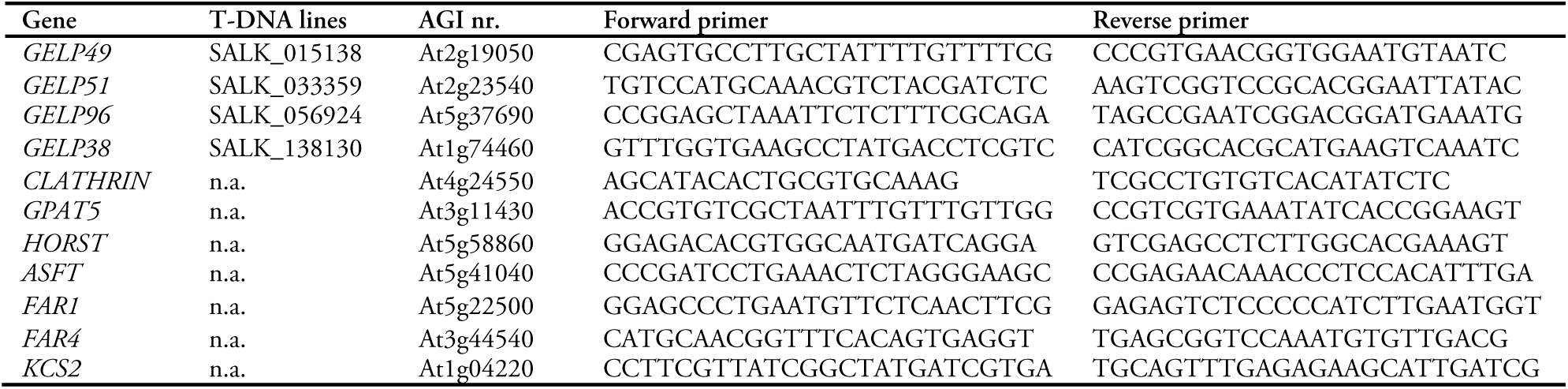
qPCR primers.

**Table S7.**
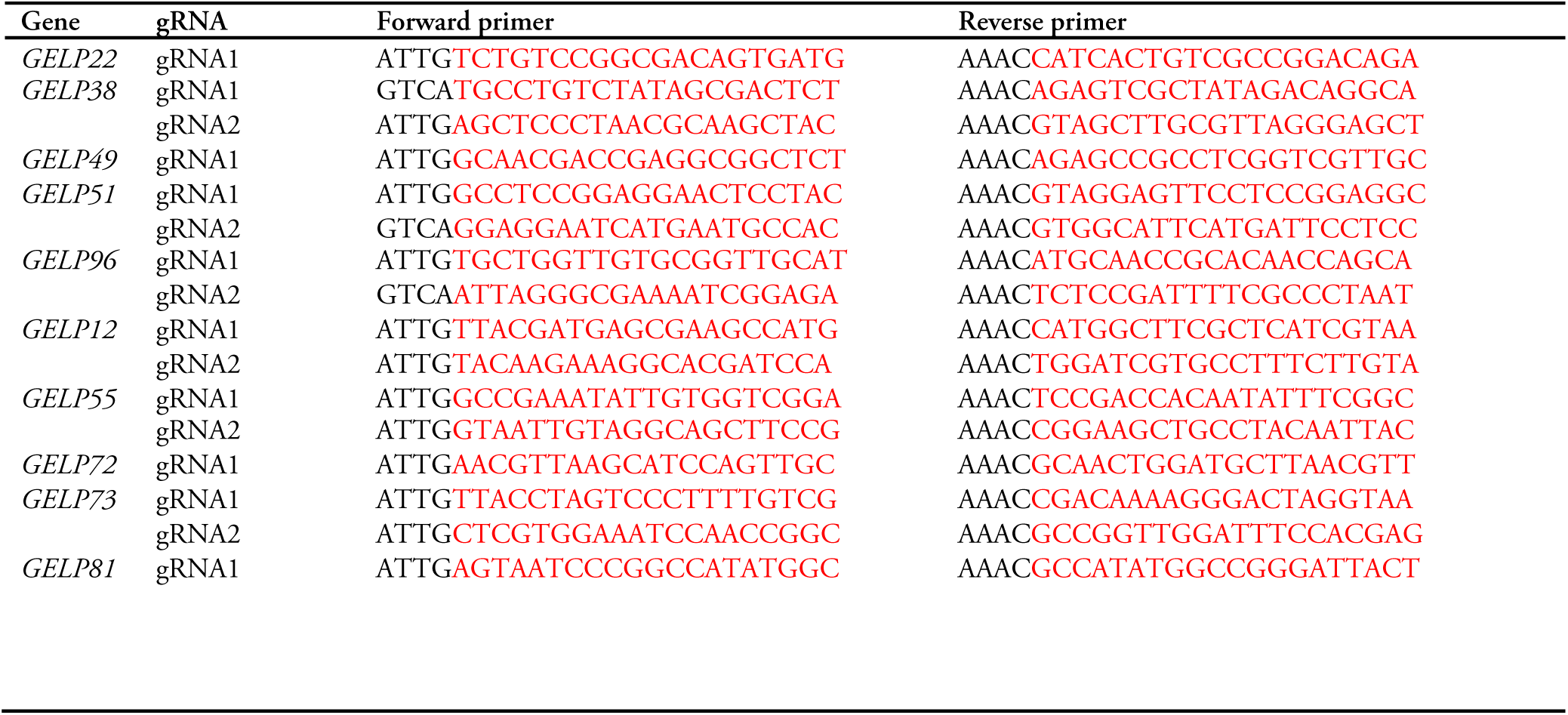
Primers for gRNA cloning (red color indicates 20 nt protospacer sequence).

## REFERENCES

Alexa, A., and Rahnenfuhrer, J. (2019). topGO: Enrichment Analysis for Gene Ontology. R package version 2381.

Anders, S., Pyl, P.T., and Huber, W. (2015). HTSeq--a Python framework to work with high-throughput sequencing data. Bioinformatics 31, 166–169.

Andersen, T.G., Molina, D., Kilian, J., Franke, R., Ragni, L., and Geldner, N. (2020). Tissue-autonomous phenylpropanoid production is essential for establishment of root barriers. bioRxiv.

Andersen, T.G., Naseer, S., Ursache, R., Wybouw, B., Smet, W., De Rybel, B., Vermeer, J.E.M., and Geldner, N. (2018). Diffusible repression of cytokinin signalling produces endodermal symmetry and passage cells. Nature 555, 529–533.

Bakan, B., and Marion, D. (2017). Assembly of the Cutin Polyester: From Cells to Extracellular Cell Walls. Plants (Basel) 6.

Banda, J., Bellande, K., von Wangenheim, D., Goh, T., Guyomarc’h, S., Laplaze, L., and Bennett, M.J. (2019). Lateral Root Formation in Arabidopsis: A Well-Ordered LRexit. Trends Plant Sci 24, 826–839.

Barberon, M., Vermeer, J.E., De Bellis, D., Wang, P., Naseer, S., Andersen, T.G., Humbel, B.M., Nawrath, C., Takano, J., Salt, D.E., et al. (2016). Adaptation of Root Function by Nutrient-Induced Plasticity of Endodermal Differentiation. Cell 164, 447–459.

Beisson, F., Li, Y., Bonaventure, G., Pollard, M., and Ohlrogge, J.B. (2007). The acyltransferase GPAT5 is required for the synthesis of suberin in seed coat and root of Arabidopsis. Plant Cell 19, 351–368.

Berhin, A., de Bellis, D., Franke, R.B., Buono, R.A., Nowack, M.K., and Nawrath, C. (2019). The Root Cap Cuticle: A Cell Wall Structure for Seedling Establishment and Lateral Root Formation. Cell 176, 1367–1378 e1368.

Clough, S.J., and Bent, A.F. (1998). Floral dip: a simplified method for Agrobacterium-mediated transformation of Arabidopsis thaliana. Plant J 16, 735–743.

Cohen, H., Fedyuk, V., Wang, C., Wu, S., and Aharoni, A. (2020). SUBERMAN regulates developmental suberization of the Arabidopsis root endodermis. Plant J.

Dobin, A., Davis, C.A., Schlesinger, F., Drenkow, J., Zaleski, C., Jha, S., Batut, P., Chaisson, M., and Gingeras, T.R. (2013). STAR: ultrafast universal RNA-seq aligner. Bioinformatics 29, 15–21.

Edqvist, J., Blomqvist, K., Nieuwland, J., and Salminen, T.A. (2018). Plant lipid transfer proteins: are we finally closing in on the roles of these enigmatic proteins? J Lipid Res 59, 1374–1382.

Fich, E.A., Segerson, N.A., and Rose, J.K. (2016). The Plant Polyester Cutin: Biosynthesis, Structure, and Biological Roles. Annu Rev Plant Biol 67, 207–233.

Fujita, S., De Bellis, D., Edel, K.H., Koster, P., Andersen, T.G., Schmid-Siegert, E., Denervaud Tendon, V., Pfister, A., Marhavy, P., Ursache, R., et al. (2020). SCHENGEN receptor module drives localized ROS production and lignification in plant roots. EMBO J, e103894.

Fukaki, H., Tameda, S., Masuda, H., and Tasaka, M. (2002). Lateral root formation is blocked by a gain-of-function mutation in the SOLITARY-ROOT/IAA14 gene of Arabidopsis. The Plant journal : for cell and molecular biology 29, 153–168.

Gasperini, D., Chetelat, A., Acosta, I.F., Goossens, J., Pauwels, L., Goossens, A., Dreos, R., Alfonso, E., and Farmer, E.E. (2015). Multilayered Organization of Jasmonate Signalling in the Regulation of Root Growth. PLoS genetics 11, e1005300.

Girard, A.L., Mounet, F., Lemaire-Chamley, M., Gaillard, C., Elmorjani, K., Vivancos, J., Runavot, J.L., Quemener, B., Petit, J., Germain, V., et al. (2012). Tomato GDSL1 is required for cutin deposition in the fruit cuticle. Plant Cell 24, 3119–3134.

Gu, Z., Eils, R., and Schlesner, M. (2016). Complex heatmaps reveal patterns and correlations in multidimensional genomic data. Bioinformatics 32, 2847–2849.

Jan, M., Gobet, N., Diessler, S., Franken, P., and Xenarios, I. (2019). A multi-omics digital research object for the genetics of sleep regulation. Sci Data 6, 258.

Kosma, D.K., Murmu, J., Razeq, F.M., Santos, P., Bourgault, R., Molina, I., and Rowland, O. (2014). AtMYB41 activates ectopic suberin synthesis and assembly in multiple plant species and cell types. Plant J 80, 216–229.

Kremer, J.R., Mastronarde, D.N., and McIntosh, J.R. (1996). Computer visualization of three-dimensional image data using IMOD. J Struct Biol 116, 71–76.

Kumpf, R.P., Shi, C.L., Larrieu, A., Sto, I.M., Butenko, M.A., Peret, B., Riiser, E.S., Bennett, M.J., and Aalen, R.B. (2013). Floral organ abscission peptide IDA and its HAE/HSL2 receptors control cell separation during lateral root emergence. Proceedings of the National Academy of Sciences of the United States of America 110, 5235–5240.

Lai, C.P., Huang, L.M., Chen, L.O., Chan, M.T., and Shaw, J.F. (2017). Genome-wide analysis of GDSL-type esterases/lipases in Arabidopsis. Plant Mol Biol 95, 181–197.

Lewis, D.R., Olex, A.L., Lundy, S.R., Turkett, W.H., Fetrow, J.S., and Muday, G.K. (2013). A Kinetic Analysis of the Auxin Transcriptome Reveals Cell Wall Remodeling Proteins That Modulate Lateral Root Development in Arabidopsis. The Plant cell.

Li, B., and Dewey, C.N. (2011). RSEM: accurate transcript quantification from RNA-Seq data with or without a reference genome. BMC Bioinformatics 12, 323.

Li, B., Kamiya, T., Kalmbach, L., Yamagami, M., Yamaguchi, K., Shigenobu, S., Sawa, S., Danku, J.M., Salt, D.E., Geldner, N., et al. (2017). Role of LOTR1 in Nutrient Transport through Organization of Spatial Distribution of Root Endodermal Barriers. Curr Biol 27, 758–765.

Li-Beisson, Y., Shorrosh, B., Beisson, F., Andersson, M.X., Arondel, V., Bates, P.D., Baud, S., Bird, D., Debono, A., Durrett, T.P., et al. (2013). Acyl-lipid metabolism. The Arabidopsis book / American Society of Plant Biologists 11, e0161.

Lucas, M., Godin, C., Jay-Allemand, C., and Laplaze, L. (2008). Auxin fluxes in the root apex co-regulate gravitropism and lateral root initiation. J Exp Bot 59, 55–66.

Naseer, S., Lee, Y., Lapierre, C., Franke, R., Nawrath, C., and Geldner, N. (2012). Casparian strip diffusion barrier in Arabidopsis is made of a lignin polymer without suberin. Proceedings of the National Academy of Sciences of the United States of America 109, 10101–10106.

Péret, B., Li, G., Zhao, J., Band, L.R., Voß, U., Postaire, O., Luu, D.-T., Da Ines, O., Casimiro, I., Lucas, M., et al. (2012). Auxin regulates aquaporin function to facilitate lateral root emergence. Nat Cell Biol 14, 991–998.

Philippe, G., Gaillard, C., Petit, J., Geneix, N., Dalgalarrondo, M., Bres, C., Mauxion, J.P., Franke, R., Rothan, C., Schreiber, L., et al. (2016). Ester Cross-Link Profiling of the Cutin Polymer of Wild-Type and Cutin Synthase Tomato Mutants Highlights Different Mechanisms of Polymerization. Plant Physiol 170, 807–820.

Philippe, G., Sorensen, I., Jiao, C., Sun, X., Fei, Z., Domozych, D.S., and Rose, J.K. (2020). Cutin and suberin: assembly and origins of specialized lipidic cell wall scaffolds. Curr Opin Plant Biol 55, 11–20.

Powers, S.K., Holehouse, A.S., Korasick, D.A., Schreiber, K.H., Clark, N.M., Jing, H., Emenecker, R., Han, S., Tycksen, E., Hwang, I., et al. (2019). Nucleo-cytoplasmic Partitioning of ARF Proteins Controls Auxin Responses in Arabidopsis thaliana. Mol Cell 76, 177–190 e175.

Salminen, T.A., Eklund, D.M., Joly, V., Blomqvist, K., Matton, D.P., and Edqvist, J. (2018). Deciphering the Evolution and Development of the Cuticle by Studying Lipid Transfer Proteins in Mosses and Liverworts. Plants (Basel) 7.

Schindelin, J., Arganda-Carreras, I., Frise, E., Kaynig, V., Longair, M., Pietzsch, T., Preibisch, S., Rueden, C., Saalfeld, S., Schmid, B., et al. (2012). Fiji: an open-source platform for biological-image analysis. Nat Methods 9, 676–682.

Schlereth, A., Moller, B., Liu, W., Kientz, M., Flipse, J., Rademacher, E.H., Schmid, M., Jurgens, G., and Weijers, D. (2010). MONOPTEROS controls embryonic root initiation by regulating a mobile transcription factor. Nature 464, 913–916.

Shi, J.X., Malitsky, S., De Oliveira, S., Branigan, C., Franke, R.B., Schreiber, L., and Aharoni, A. (2011). SHINE transcription factors act redundantly to pattern the archetypal surface of Arabidopsis flower organs. PLoS genetics 7, e1001388.

Siligato, R., Wang, X., Yadav, S.R., Lehesranta, S., Ma, G., Ursache, R., Sevilem, I., Zhang, J., Gorte, M., Prasad, K., et al. (2016). MultiSite Gateway-Compatible Cell Type-Specific Gene-Inducible System for Plants. Plant Physiol 170, 627–641.

Stoeckle, D., Thellmann, M., and Vermeer, J.E. (2018). Breakout-lateral root emergence in Arabidopsis thaliana. Curr Opin Plant Biol 41, 67–72.

Swarup, K., Benkova, E., Swarup, R., Casimiro, I., Peret, B., Yang, Y., Parry, G., Nielsen, E., De Smet, I., Vanneste, S., et al. (2008). The auxin influx carrier LAX3 promotes lateral root emergence. Nat Cell Biol 10, 946–954.

Tian, Q., Uhlir, N.J., and Reed, J.W. (2002). Arabidopsis SHY2/ IAA3 inhibits auxin-regulated gene expression. The Plant cell 14, 301–319.

Ursache, R., Andersen, T.G., Marhavy, P., and Geldner, N. (2018). A protocol for combining fluorescent proteins with histological stains for diverse cell wall components. Plant J 93, 399–412.

Vermeer, J.E., von Wangenheim, D., Barberon, M., Lee, Y., Stelzer, E.H., Maizel, A., and Geldner, N. (2014). A spatial accommodation by neighboring cells is required for organ initiation in Arabidopsis. Science 343, 178–183.

Vishwanath, S.J., Delude, C., Domergue, F., and Rowland, O. (2015). Suberin: biosynthesis, regulation, and polymer assembly of a protective extracellular barrier. Plant cell reports 34, 573–586.

Voß, U., Wilson, M.H., Kenobi, K., Gould, P.D., Robertson, F.C., Peer, W.A., Lucas, M., Swarup, K., Casimiro, I., Holman, T.J., et al. (2015). The circadian clock rephases during lateral root organ initiation in Arabidopsis thaliana. Nature communications 6, 7641.

Yadav, V., Molina, I., Ranathunge, K., Castillo, I.Q., Rothstein, S.J., and Reed, J.W. (2014). ABCG transporters are required for suberin and pollen wall extracellular barriers in Arabidopsis. Plant Cell 26, 3569–3588.

Yeats, T.H., Martin, L.B., Viart, H.M., Isaacson, T., He, Y., Zhao, L., Matas, A.J., Buda, G.J., Domozych, D.S., Clausen, M.H., et al. (2012). The identification of cutin synthase: formation of the plant polyester cutin. Nat Chem Biol 8, 609–611.

